# Circadian oscillation in primary cilium length by clock genes regulate fibroblast cell migration

**DOI:** 10.1101/2023.01.24.525311

**Authors:** Ryota Nakazato, Yuki Matsuda, Faryal Ijaz, Koji Ikegami

## Abstract

Various mammalian cells have autonomous cellular clocks that are produced by the transcriptional cycle of clock genes. Cellular clocks provide circadian rhythms for cellular functions via transcriptional and cytoskeletal regulation. The vast majority of mammalian cells possess a primary cilium, an organelle protruding from the cell surface. Here, we investigated the little-known relationship between circadian rhythm and primary cilia. The length and number of primary cilia showed circadian dynamics both in vitro and in vivo. The circadian rhythm of primary cilium length was abolished by SR9011 and Bmal1 knockout. A centrosomal protein, pericentrin, transiently accumulates in centriolar satellites, the base of primary cilia at the shortest cilia phase, and induces elongation of primary cilia at the longest cilia phase in the circadian rhythm of primary cilia. In addition, rhythmic cell migration during wound healing depends on the length of primary cilia and affects the rate of wound healing. Our findings demonstrate that the circadian dynamics of primary cilia length by clock genes control fibroblast migration and could provide new insights into chronobiology.

## Introduction

Circadian rhythmicity is observed in various biological phenomena in organisms, including the sleep/wake cycle, ranging from bacteria to mammals. This system adjusts physiological processes to an approximately 24-h cycle, and adapts it to the external environment of the day–night cycle. The suprachiasmatic nucleus (SCN) of hypothalamus is believed to be a “central clock” that synchronizes the clock in every cell and tissues in vivo (Yamazaki *et al*, 2000). Disruption of the central clock causes various diseases, including abnormal sleep behaviors, obesity, and vascular diseases (Nakazato *et al*, 2017; Turek *et al*, 2005; Zee & Vitiello, 2009). Circadian rhythms are generated by a transcriptional feedback loop of “clock gene” expression. The core loop consisted of two positive and two negative regulators. The brain and muscle aryl hydrocarbon receptor nuclear translocator-like protein-1 (BMAL1; also known as ARNTL) and circadian locomotor output cycles protein kaput (CLOCK) function as two positive regulators (Gekakis *et al*, 1998). The BMAL1/CLOCK complex binds to the E-box motif and drives the transcription of various genes, including period (PER) and cryptochrome (CRY), which inhibit the transcriptional activity of the BMAL1/CLOCK complex as negative regulators of the loop (Griffin *et al*, 1999; Yamaguchi *et al*, 2000). This core feedback loop, which takes place in a 24-h cycle generates circadian rhythmicity (Kume *et al*, 1999).

Most organs, tissues, and cells in the non-SCN also have an intrinsic circadian clock, while the center of the circadian clock is the SCN (Shimba *et al*, 2005; Storch *et al*, 2002; Young *et al*, 2001). Negative feedback loops by clock genes exist in individual cells, and oscillations are observed in each single cell. The oscillations are also called an autonomous cellular clock. The autonomous cellular clock provides circadian rhythms for cell-specific functions in an SCN-independent manner. This clock regulates the cytoskeleton and organelles, and provides rhythmic changes in cellular functions (Mofatteh *et al*, 2021). For example, the autonomous cellular clock regulates fibroblast migration by controlling actin dynamics (Hoyle *et al*, 2017). Human studies have shown that burns sustained at night take longer to heal than those sustained during the day, possibly because fibroblast migration to the wound is regulated by clock genes (Hoyle *et al*., 2017). The accumulation of key mitochondrial enzymes is regulated by Per1/2 and exhibits diurnal oscillations, which are thought to adapt mitochondrial activities to changes in body activity and rest, feeding, and fasting throughout the day (Neufeld-Cohen *et al*, 2016). The formation of daughter centrioles has been reported to exhibit diurnal oscillations due to Plk4 oscillations in centrioles and centriolar duplicating independent of the cell cycle (Aydogan *et al*, 2020).

Most cells possess a primary cilium, which is a small organelle protruding from the cell surface that functions as a sensory antenna for cells to detect extracellular signals (Phua *et al*, 2015; Singla & Reiter, 2006). The primary cilium is recognized as a crucial organelle for various cellular processes including proliferation, differentiation, and migration (Ezratty *et al*, 2011; Higginbotham *et al*, 2012; Irigoín & Badano, 2011; Rohatgi *et al*, 2007). Defects in primary cilia due to congenital factors such as genetic mutations in *BBS*, *PKD* and *OFD*, induce several diseases “ciliopathy”, including obesity, polycystic kidney, and retinopathy (Quinlan *et al*, 2008; Reiter & Leroux, 2017). Primary cilia are immotile. In the cell cycle, primary cilia assemble in non-dividing cells in the G_0_/G_1_ phase and disassemble with cell cycle entry (Plotnikova *et al*, 2009). We previously reported that the decapitation of primary cilia and release of vesicles are signals that induce primary cilia disassembly and drive the cell cycle (Phua *et al*, 2017).

Morphological dynamics in primary cilia (including length changes) can also contribute to the regulation of cellular functions and extracellular signal reception such as fluid flow, Hedgehog (Hh) and Wnt signals (Broekhuis *et al*, 2013; Keeling *et al*, 2016). Cell sensitivity to mechanical stimuli increases depending on the length of primary cilia (Spasic & Jacobs, 2017). Aldosterone-stimulated increase in cell size of renal collecting duct epithelial cells seems to be dependent on the length of primary cilia (Komarynets *et al*, 2020). Despite of accumulation of these evidence, the mechanisms regulating primary cilium morphology and the physiological significance of alterations in primary cilium morphology have not been fully elucidated. Here, we investigated the relationship between the circadian rhythm and primary cilia. Recently, it was reported that the length of primary cilia in the SCN exhibits circadian rhythms and could contribute to the maintenance of endogenous circadian oscillations (Tu *et al*, 2023). We demonstrated that the length of primary cilia in cultured murine fibroblasts, neurons, and astrocytes of the murine brain exhibits circadian dynamics similar to the rhythmic oscillations of clock genes. Wound healing assay revealed that the circadian dynamics of primary cilium length regulate fibroblast migration. Our data indicate that circadian rhythms in primary cilium length produced by clock genes regulate cell migration. The findings provide new insights into chronobiology.

## Results

### Ciliogenesis and primary cilium length in cultured cells exhibit circadian rhythms

To investigate the relationship between primary cilia and circadian rhythm, we analyzed changes in primary cilium length and clock gene expression in NIH/3T3 mouse embryonic fibroblasts. In cultured cells, primary cilia assembly is induced by serum starvation-induced cell cycle arrest (R W Tucker, 1979). Cultured cells also have autonomous clock gene expression rhythms, but their individual phases are disparate (Nagoshi *et al*, 2004). Dexamethasone (DEX), a synthetic corticosteroid, induces Per1 expression and synchronizes the clock between cells (Nagoshi *et al*., 2004). In this study, ciliogenesis was induced by incubation in culture medium for 24 h with serum concentration decreased from 10% to 1%, followed by 2-h exposure to DEX to synchronize clock gene expression (Fig. 1A). The DEX-induced synchronization of clock genes expression was verified by using Per1::Luc-expressing NIH/3T3 cells. The cells were exposed to DEX for 2 h to synchronize the expression of clock genes. Bioluminescence was measured every 15 min for 96 h using photomultiplier tube detector assemblies. The oscillatory pattern of luciferase activity was observed in cells exposed to DEX (Fig. 1B). Based on these results, we used this synchronization regimen in subsequent experiments.

**Figure 1.**
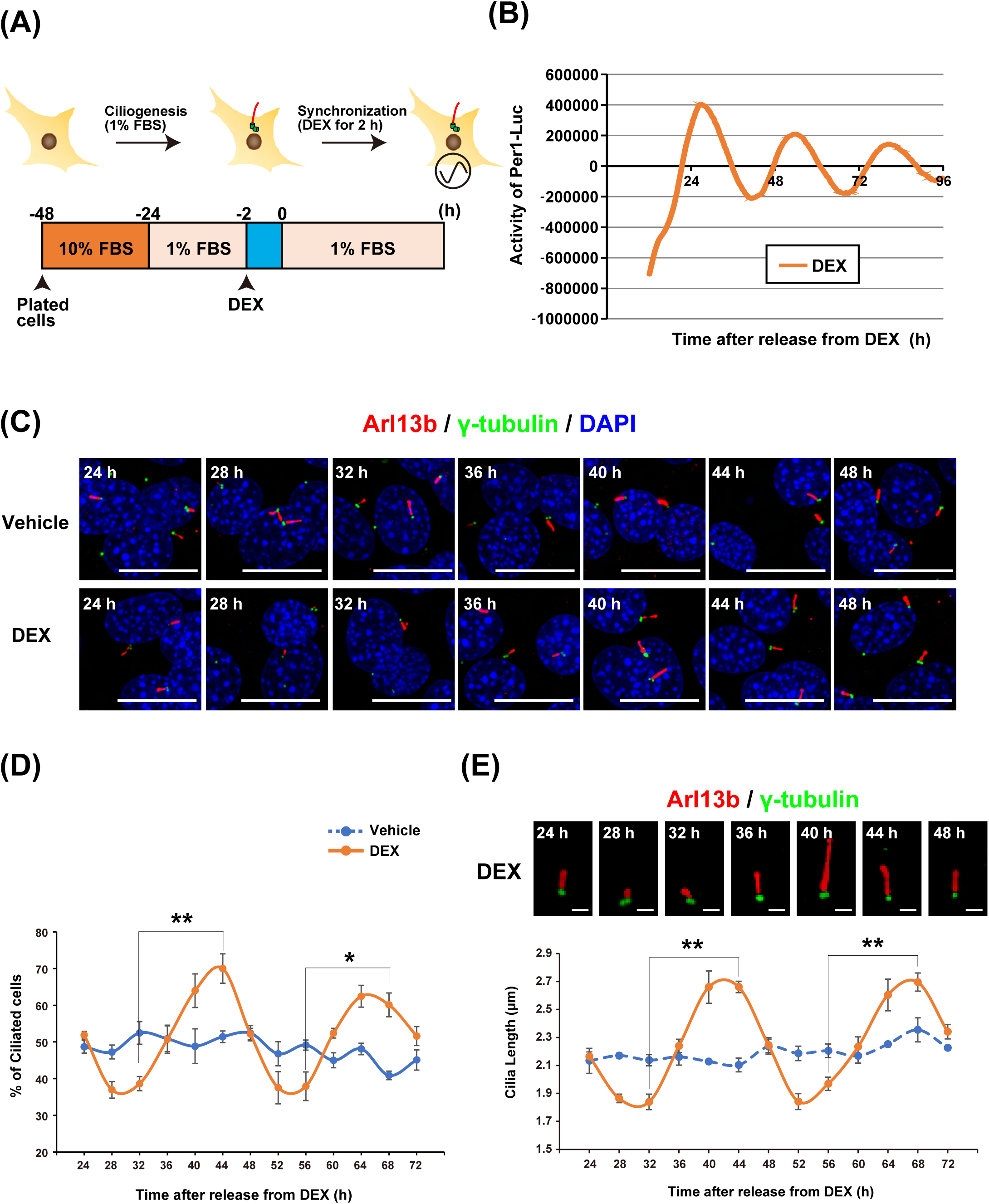
Length of primary cilia in cultured cells exhibits circadian rhythm. A) Schematic diagram of the protocol for induction of ciliogenesis by serum starvation and synchronization of clock gene expression by exposure of cultured NIH/3T3 cells to dexamethasone (DEX). B) Real-time luciferase bioluminescence assay for Per1 promoter activity levels in Per1::Luc-expressing NIH/3T3 cells exposed to DEX as in A. C) Immunostaining for primary cilia in NIH/3T3 cells at the indicated time points following treatment with the protocol used in panel A. Red denotes labeling of cilia by anti-Arl13b antibody, green denotes labeling of centrosomes by anti-γ-tubulin antibody, and blue denotes labeling of DNA by DAPI. White arrows point to primary cilia. Scale bar, 20 µm. D) Quantitative analysis of the percentage of Arl13b^+^ NIH/3T3 cells at each indicated time point. The data are from three independent experiments. E) Quantitative analysis of the primary cilium length in NIH/3T3 cells at each indicated time point. The data are from three independent experiments. Representative views of DEX-synchronized primary cilia are provided on the graph. Scale bar, 2 µm. Data information: Data in panels D and E are presented as mean ± SEM. *P≤0.05 and **P≤0.01 (One-way ANOVA).

To investigate the influence of the cell intrinsic clock on primary cilia, we performed immunostaining of primary cilia in synchronized NIH/3T3 cells. NIH/3T3 cells were synchronized by DEX, then fixed at 24, 28, 32, 36, 40, 44, 48, 52, 56, 60, 64, 68 and 72 h after switching the medium to DEX-free one. Primary cilia were visualized by staining Arl13b, a marker of primary cilia, and the base of primary cilia by staining γ-tubulin, a peri-centrosome component (Fig. 1C). In NIH/3T3 cells treated with vehicle, where conditions were unsynchronized, Arl13b-positive primary cilia were observed in approximately 50% of the total cells at all times (Fig. 1C, 1D). In DEX-treated synchronized conditions, cells with Arl13b-positive primary cilia were minimum 28 to 32 h and 52 to 54 h after synchronization (36.93 ± 2.24% at 28 h, 38.61 ± 1.91% at 32 h, 37.52 ± 4.39% at 52 h, 37.92 ± 3.90% at 56 h, Fig. 1D), and maximum 40 to 44 h and 64 to 68 h after synchronization (63.97 ± 4.58% at 40 h, 70.00 ± 4.01% at 44 h, 62.45 ± 2.97% at 64 h, 60.11 ± 3.24% at 68 h, Fig. 1D). At other time points, the proportion of ciliated cells was approximately 50% (Fig. 1D). We further examined the effect of the circadian rhythms on the length of primary cilia. The average length of the primary cilia in unsynchronized cells was approximately 2 µm (Fig. 1E). Primary cilia length in synchronized cells was shortest 28 to 32 h and 52 to 56 h after synchronization (1.86 ± 0.03 µm at 28 h, 1.84 ± 0.06 µm at 32 h, 1.84 ± 0.06 µm at 52 h and 1.97 ± 0.05 µm at 56 h, Fig. 1E) and longest 40 to 42 h and 64 to 68 h after synchronization (2.66 ± 0.12 µm at 40 h, 2.66 ± 0.04 µm at 44 h, 2.60 ± 0.11 µm at 64 h and 2.70 ± 0.06 µm at 68 h, Fig. 1E). At other time points, the length of primary cilia was approximately 2 µm (Fig. 1E). To monitor primary cilia in live cells, we performed time-lapse imaging using NIH/3T3 cells that expressed Arl13b-venus under controls by endogenous Arl13b promoter. The length of individual primary cilia showed obvious oscillation with ∼24 h intervals in live-cell imaging (Movie EV3, EV4 and Fig. EV1A). These data indicate that both the number and length of primary cilia exhibit circadian oscillations.

### Circadian changes of primary cilium length is regulated by clock genes

To examine whether the circadian oscillation of primary cilia length is mediated by clock gene expression, NIH/3T3 cells synchronized by DEX treatment were exposed to SR9011, a synthetic agonist for Rev-Erb, which suppresses Bmal1 and Clock expression (Sulli *et al*, 2018) (Fig. 2A). SR9011 also inhibits autophagy and lipogenesis in cancer cells at 20 µM (Sulli *et al*, 2018). The blocking of autophagy is reported to cause primary cilia elongation (Pampliega & Cuervo, 2016; Pampliega *et al*, 2013). Indeed, 10 µM SR9011 induced the elongation of primary cilia (average of primary cilium length 4.17 ± 0.18 µm for 10 µM SR9011 versus 2.81 ± 0.20 µm for control) in unsynchronized cells (Fig. 2B and 2C). We thus sought out an appropriate concentration of SR9011 that did not induce primary cilia elongation, but inhibited the circadian dynamics of primary cilia. To this end, we treated NIH/3T3 cells with 2-to-10-µM SR9011 at 30 h after synchronization, at which the shortest cilia length and maximal Bmal/Clock activity were observed. Total RNA was collected at 30 h after 2-h exposure to DEX, cDNA was synthesized by reverse transcription, and the expression level of the clock gene Bmal1 mRNA (gene symbol is *Arntl*) was measured. Two micromolar SR9011 was enough to significantly suppress Bmal1 expression (Fig. 2D).

**Figure 2.**
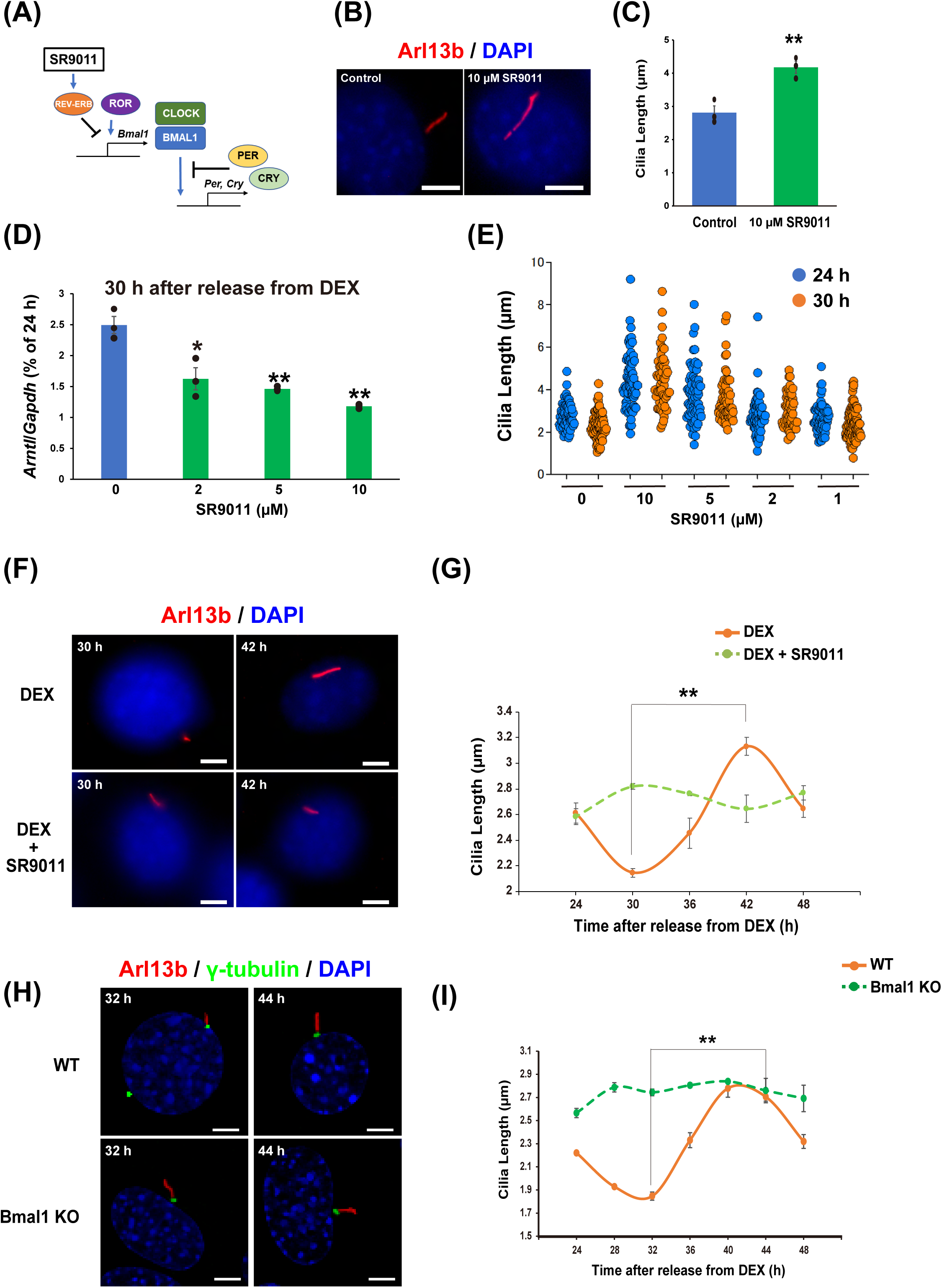
Clock genes contribute to circadian rhythms of primary cilium length. A) Schematic diagram of molecular core clock mechanism and SR9011 pharmacology. B) Immunostaining of primary cilia in NIH/3T3 cells treated with 10 µM SR9011. Scale bar, 5 µm. C) Quantitative analysis of length of primary cilia in panel B. The data are from three independent experiments. D) Real-time qPCR analysis of Bmal1 mRNA expression levels in the NIH/3T3 cells incubated with SR9011 at 30 h after release from DEX. The data are from three independent experiments. E) Quantitative analysis of primary cilium length in NIH/3T3 cells treated with the indicated concentration of SR9011 24 or 30 h after release from DEX, as in panel A. The data are from over 50 cells per condition. F) Immunostaining for primary cilia in NIH/3T3 cells treated with 2 µM SR9011 30 or 42 h after release from DEX. Scale bar, 5 µm. G) Quantitative analysis of primary cilium length in NIH/3T3 cells at each indicated time point after release from DEX in panel F. The data are from three independent experiments. H) Immunostaining for primary cilia in WT or Bmal1-KO NIH/3T3 cells at indicated time point after release from DEX. Scale bar, 5 µm. I) Quantitative analysis of primary cilium length in WT or Bmal1-KO NIH/3T3 cells at each indicated time point after rel ease from DEX in panel H. The data are from three independent experiments. Data information: Data in panels C, D, G and I are presented as mean ± SEM. *P≤0.05 and **P≤0.01 (One-way ANOVA).

SR9011 at 2 µM did not induce the abnormal primary cilia elongation at 24 h after synchronization (Fig. 2E). In contrast, the primary cilia shortening at 30 h after synchronization was canceled by the same concentration of SR9011 (average primary cilia length 2.15 ± 0.03 µm with DEX versus 2.82 ± 0.02 µm with DEX and SR9011) (Fig. 2E). At 1 µM, SR9011 failed to block the shortening of primary cilia 30 h after synchronization (Fig. 2E). We then examined whether the circadian oscillation of primary cilia length was abolished with SR9011 at this concentration. The elongation of primary cilia at 42 h after synchronization was also abolished by 2 µM SR9011 (3.13 ± 0.07 µm with DEX versus 2.64 ± 0.11 µm with DEX and SR9011) (Fig. 2F and 2G). Hence, SR9011 significantly suppressed the circadian oscillation of primary cilium length in NIH/3T3 cells (Fig. 2G). This suppression continues for at least 72 h after synchronization by DEX (Fig. EV1B).

To examine direct effects of clock genes on primary cilium length, we generated Bmal1-KO NIH/3T3 cells using CRISPR/Cas9 and measured the length of primary cilia. The rhythm of primary cilium length observed in wildtype (WT) cells is abolished in Bmal1-KO NIH/3T3 cells (2.74 ± 0.03 µm at 32 h and 2.75 ± 0.11 µm at 44 h in Bmal1 KO NIH/3T3 cells) (Fig. 2H and 2I). These results indicate that the circadian oscillation of primary cilia length is mediated by rhythmic expression of clock genes.

### Primary cilium length in murine brain also exhibits circadian rhythms

Next, to assess whether primary cilium length also shows circadian dynamics *in vivo*, we performed immunostaining of murine brains collected at different time points of day and night. In the adult murine brain, marker proteins for primary cilia differ among cell types (Sterpka & Chen, 2018). Hippocampal dentate gyrus of the murine brain was used for immunohistochemistry targeting adenylate cyclase 3 (AC3) as a primary ciliary neuronal marker, NeuN (neuronal marker), glial fibrillary acidic protein (GFAP, astroglial marker), and Arl13b (Fig. 3A). AC3-positive primary cilia were observed in NeuN-positive cells but not in GFAP-positive cells (Fig. 3A; left and middle panels). Arl13b-positive primary cilia were observed in GFAP-positive cells (Fig. 3A; right panel). Perfusion fixation was performed in adult mice using 4% paraformaldehyde (PFA) at Zeitgeber Time (ZT)5 and ZT17, followed by immunohistochemical staining. The length of primary cilia in neurons in the murine hippocampus was significantly longer at ZT17 than at ZT5 (Fig. 3B and 3C). The primary cilia length in astrocytes was also markedly longer at ZT17 than at ZT5 (Fig. 3D and 3E). These results indicate that primary cilium length *in vivo* also exhibits circadian oscillations.

**Figure 3.**
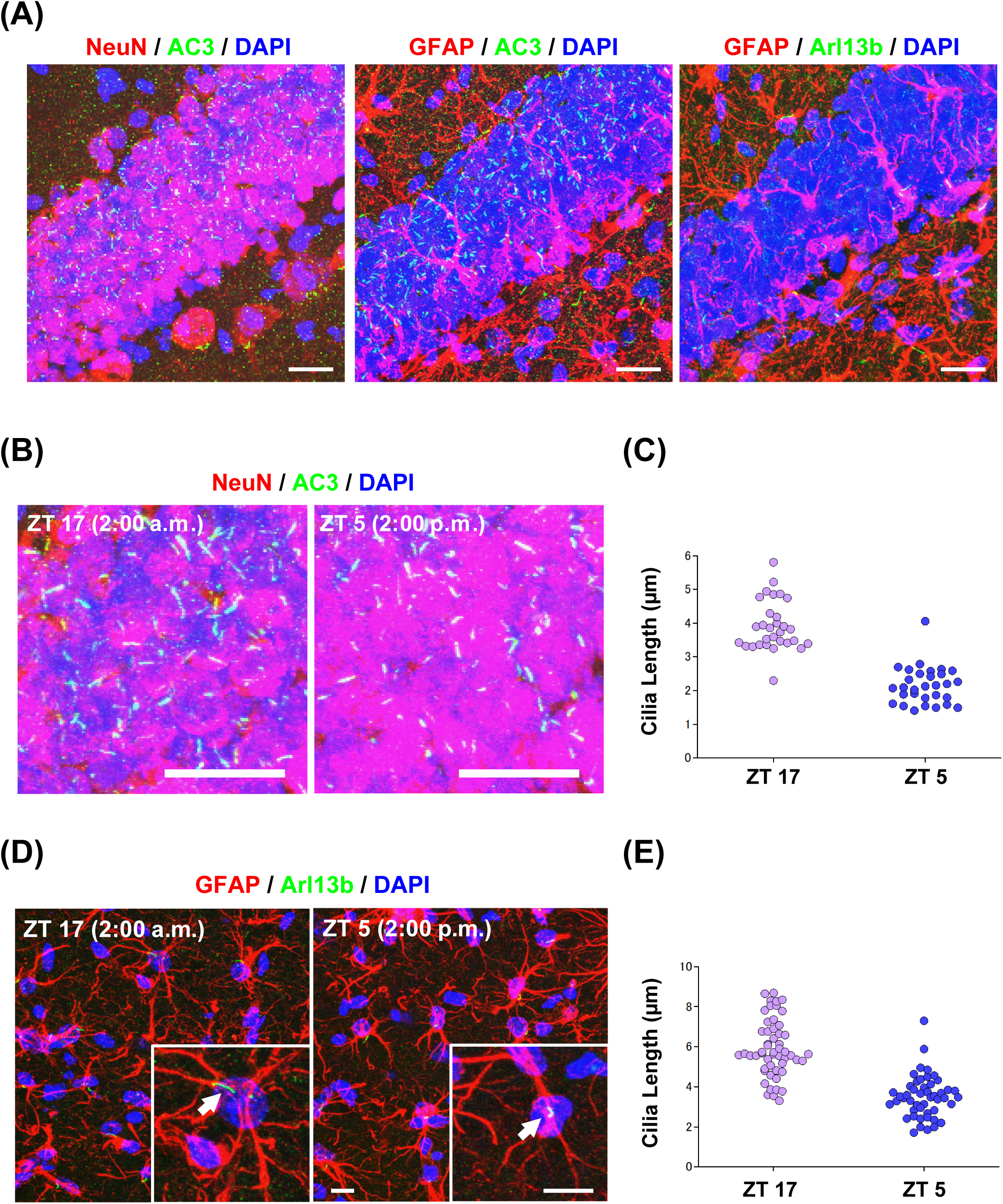
Length of primary cilia also exhibits circadian rhythm in murine brain. A) Immunostaining for primary cilia on neurons (green; labeled by anti-AC3 antibody) and neural markers (red; labeled with anti-NeuN antibody) or primary cilia on astrocytes (green; labeled by anti-Arl13b antibody) and astroglial markers (red; labeled by anti-GFAP antibody) in murine brain hippocampus. Scale bar, 20 µm. B) Immunostaining for primary cilia in neurons in the hippocampus of perfusion-fixed murine brains at ZT5 or ZT17. Scale bar, 20 µm. C) Quantitative analysis of primary cilium length in panel B (n = 31 cilia per condition). D) Immunostaining for primary cilia on astrocytes in the hippocampus of perfusion-fixed murine brains at ZT5 or ZT17. Scale bar, 5 µm. White arrows point to primary cilia. E) Quantitative analysis of primary cilium length from panel D (n = 48-53 cilia per condition).

### Pericentrin, but not PCM1, accumulates at the periphery of the centrosome in a clock-dependent manner

To investigate the mechanism by which the cell intrinsic clock regulates the length of the primary cilia, we focused on centriolar satellites. Centriolar satellites are an array of 70-100-nm granules at the base of the primary cilia that regulate the transport of centrosomal and ciliary proteins. They are also involved in the formation of primary cilia (Odabasi *et al*, 2019; Quarantotti *et al*, 2019). Pericentrin (PCNT) is a centrosomal protein that localizes to centriolar satellites through binding to pericentriolar material 1 (PCM1), and is involved in primary cilia assembly (Jurczyk *et al*, 2004). In this study, immunostaining for PCNT was performed in NIH/3T3 cells synchronized with DEX treatment (Fig. 4A). PCNT-positive centriolar satellites transiently increased in NIH/3T3 cells 30 h after synchronization with primary cilium length minimum. The average area of PCNT-positive granules was 11.95 ± 0.55 µm^2^ 24 h, 29.59 ± 3.33 µm^2^ 30 h, 12.64 ± 0.87 µm^2^ 36 h, 12.02 ± 1.38 µm^2^ 42 h, and 13.78 ± 1.32 µm^2^ 48 h after synchronization (Fig. 4B). The increase in PCNT-positive centriolar satellites at 30 h was suppressed by 2 µM SR9011, with an average area of PCNT-positive granules of 17.49 ± 1.35 µm^2^ in cells exposed to DEX and 2 µM SR9011 (Fig. 4A and 4B). We also performed a western blotting assay using whole cell lysate of NIH/3T3 cells synchronized with DEX to test whether the oscillation of PCNT-positive granules occurs through changes in the amount of PCNT protein. No significant difference in the amount of PCNT protein was observed at any time point after synchronization (Fig. 4C). Measurement of PCNT-positive puncta revealed that PCNT-positive granules were mainly increased at a distance between 4 and 10 µm from the center of centrosome, rather than on centrosome at 30 h after synchronization (Fig. 4D). These results indicate that the transient increase in PCNT-positive centriolar satellites observed in NIH/3T3 cells 30 h after synchronization is caused by changes in subcellular localization.

**Figure 4.**
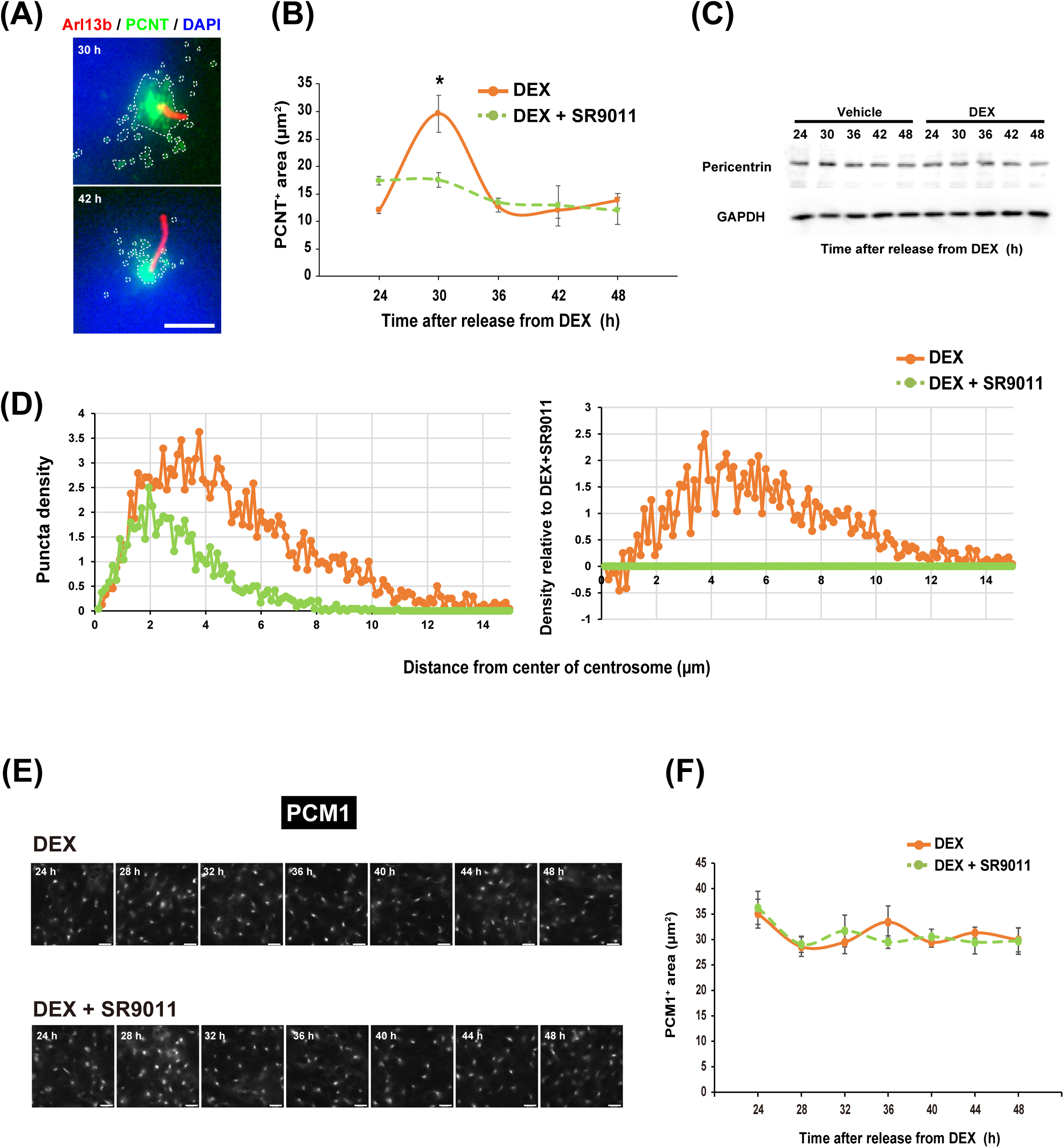
Pericentrin accumulates around the base of shortest primary cilia. A) Immunostaining for primary cilia and pericentrin. Arl13b is stained red, pericentrin (PCNT) green, and DNA blue (by DAPI) in NIH/3T3 cells at 30 and 42 h after release from DEX. Scale bar, 5 µm. White dotted lines enclose PCNT-positive granules. B) Quantitative analysis of areas positive for PCNT in NIH/3T3 cells treated with SR9011 or dimethyl sulfoxide (DMSO) as vehicle at the indicated time points after release from DEX. The data are from three independent experiments. C) Western blot analyses of expression of PCNT and GAPDH (as an internal standard) in NIH/3T3 cells at each indicated time point after release from DEX. D) Quantitative analysis of puncta density for PCNT in NIH/3T3 cells treated with SR9011 or vehicle at 30 h after release from DEX (n = 24 cells per condition). E) Immunostaining for PCM1 in NIH/3T3 cells treated with SR9011 or vehicle at indicated time points after release from DEX. Scale bar, 5 µm. F) Quantitative analysis of areas positive for PCM1 in NIH/3T3 cells treated with SR9011 or vehicle at the indicated time points after release from DEX. The data are from three independent experiments. Data information: In panels B and F, data are presented as mean ± SEM. *P≤0.05 (One-way ANOVA).

We also examined whether intracellular localization of PCM1 showed the similar pattern with PCNT after synchronization with DEX. To this end, we stained NIH/3T3 cells with anti-PCM1 antibodies. Unexpectedly, PCM1 exhibited intracellular behaviors different from those of PCNT; PCM1 did not show clear circadian oscillation (Fig. 4E and 4F). These results indicate that the transient accumulation in PCNT-positive centriolar satellites at 30 h after synchronization occurred through pathways different from PCM1-dependent ones.

### Transient increase in PCNT-positive centriolar satellites regulates circadian rhythm of primary cilium length

It has been reported that correct subcellular localization of centriolar satellites is necessary for the formation of primary cilia (Aydin *et al*, 2020). The elongation of primary cilia is known to take place over several hours (Westlake *et al*, 2011). To investigate the effect of centriolar satellites increasing at the base of primary cilia on the circadian dynamics of primary cilium length, we inhibited transport of centriolar satellites to the base of primary cilia. Centriolar satellites are transported to the base of primary cilia via microtubule transport (Dammermann & Merdes, 2002). We used colchicine, an inhibitor of microtubule polymerization, to block the accumulation of PCNT-positive granules at the base of the primary cilia. The effect of colchicine was confirmed by immunostaining the microtubules with anti-α-tubulin antibody. Microtubules visible in control NIH/3T3 cells (Fig. 5A; left panel) were disrupted by 200 nM colchicine (Fig. 5A; right panel). Colchicine inhibited the transient increase in PCNT-positive centriolar satellites at 30 h after synchronization in a concentration-dependent manner, to a level comparable to that of unsynchronized control cells (Fig. 5B and 5C). We then examined whether the blockade of PCNT-positive centriolar satellites accumulation at 30 h after synchronization impaired later primary cilia elongation. To this end, we exposed DEX-synchronized NIH/3T3 cells to 200 nM colchicine for 2 h at 28 h after synchronization, and subsequently measured the length of primary cilia visualized by immunostaining at 30 h and 42 h after synchronization (Fig. 5D). At 30 h after synchronization, the administration of colchicine made no significant difference to the length of primary cilia compared to control cells. However, at 42 h after synchronization (the longest cilia phase), 200 nM colchicine markedly suppressed the length of primary cilia compared to control cells (average primary cilia length 3.11 ± 0.10 µm for DEX versus 2.28 ± 0.09 µm for DEX with colchicine) (Fig 5E, 5F and 5G).

**Figure 5.**
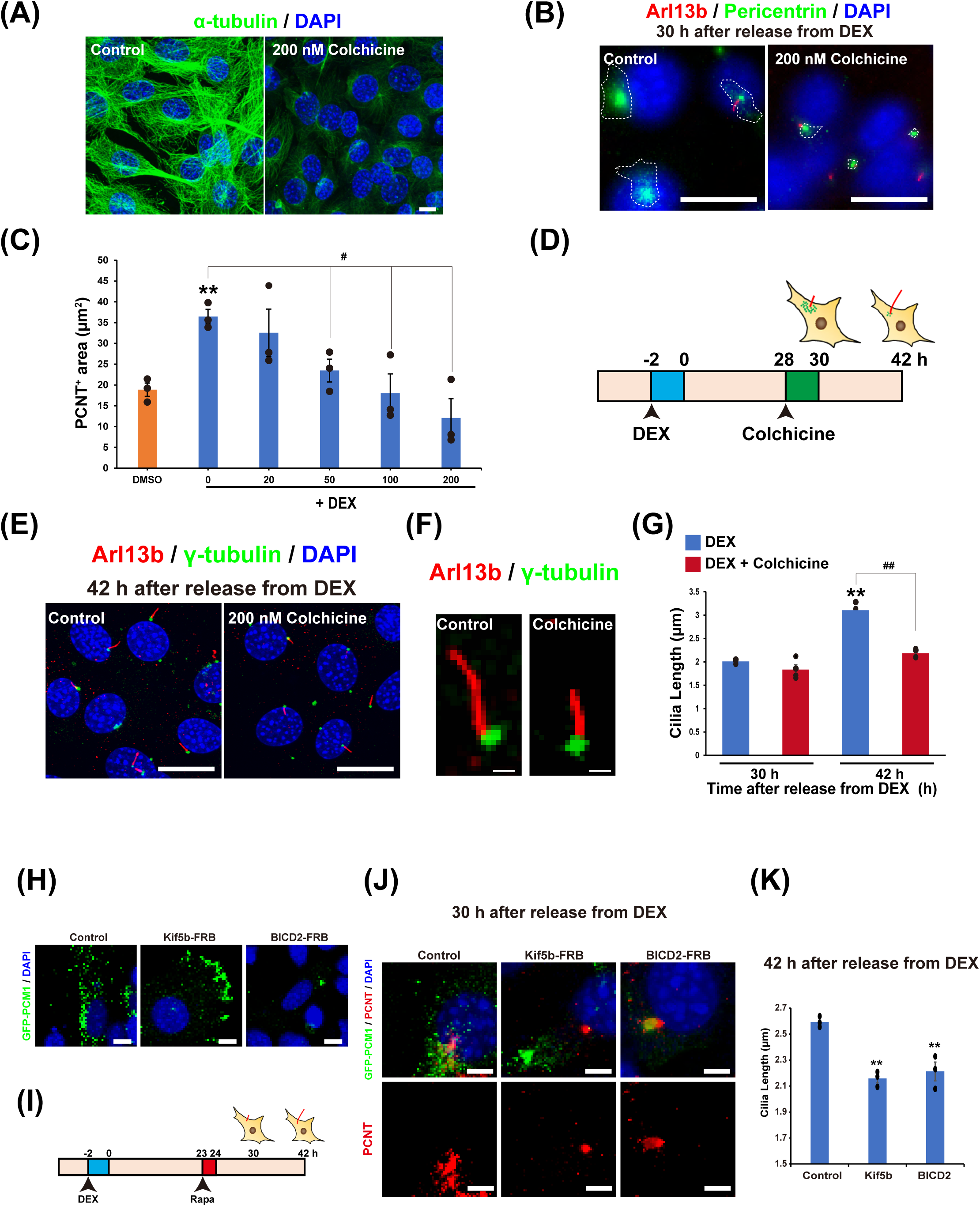
The effect of PCNT-positive centriolar satellites on circadian rhythm of primary cilium length. A) Immunostaining for microtubules (green; labeled with anti-α-tubulin antibody) and DNA (blue; DAPI) in NIH/3T3 cells incubated with 200 nM colchicine or DMSO as vehicle for 2 h. Scale bar, 10 µm. B) Immunostaining for primary cilia (red, Arl13b; green, PCNT) in NIH3T3 cells treated with 200 nM colchicine or DMSO 30 h after release from DEX. White dotted lines enclose PCNT-positive granules. Scale bar, 20 µm. C) Quantitative analysis of the areas positive for PCNT in NIH/3T3 cells with the indicated concentration of colchicine 30 h after release from DEX. The data are from three independent experiments. D) Schematic diagram of the protocol for NIH/3T3 cells synchronized with DEX and exposure to colchicine. E) Immunostaining of primary cilia (red; Arl13b) and centrosomes (green; γ-tubulin) in NIH/3T3 cells treated with 200 nM colchicine or DMSO 42 h after release from DEX. Scale bar, 20 µm. F) Representative views of primary cilia for panel I. Scale bar, 2 µm G) Quantitative analysis of primary cilium length in NIH/3T3 cells treated with 200 nM colchicine or DMSO 30 or 42 h after release from DEX. The data are from four independent experiments. H) Representative images of PCM1 redistribution to cell periphery or center upon inducible dimerization PCM1 with the plus and minus end-directed molecular motor proteins. Immunostaining for GFP in NIH/3T3 cells expressing GFP-PCM1-FKBP alone (indicated as Control), co-expressing GFP-PCM1-FKBP with HA-Kif5b-FRB (indicated as Kif5b-FRB) or HA-BICD2-FRB (indicated as BICD2-FRB). Cells were treated with rapamycin for 1 h at 23 h after release from DEX and fixed 6 h later. Scale bar, 10 µm. I) Schematic diagram of the protocol for treatment of rapamycin (Rapa) to NIH/3T3 cells exposed to DEX. J) Immunostaining for GFP (green) and PCNT (red) in NIH/3T3 cells expressing indicated plasmid at 30 h after release from DEX. Scale bar, 5 µm. K) Quantitative analysis of primary cilium length in NIH/3T3 cells expressing indicated plasmids at 42 h after release from DEX. The data are from three independent experiments. Data information: Data in panels C, G, and K are presented as mean ± SEM. **P≤0.01, #P≤0.05 and ##P≤0.01 (One-way ANOVA).

We further examined whether the transient localization of PCNT to centriolar satellites at 30 h is required for the following primary cilia elongation by perturbing PCNT localization with chemical dimerization system. We chose PCM1 as the target of chemical dimerization, since handling of PCNT itself is quite difficult due to its huge molecular size and intracellular PCNT translocation to centriolar satellites depends on PCM1 (Dammermann & Merdes, 2002). PCM1 fused to GFP at the N-terminus and FKBP at C-terminus (GFP-PCM1-FKBP), and kinesin family member 5B (Kif5b) fused to FRB at C-terminus (Kif5b-FRB) or dynein/dynactin cargo adaptor bicaudal D homolog 2 (BICD2) fused to FRB at C-terminus (BICD2-FRB) were used in the experiments (Aydin *et al*., 2020). In NIH/3T3 cells only expressing GFP-PCM1-FRB, GFP-positive particles are localized at centriolar satellites (Fig. 5H; left panel). In cells co-expressing GFP-PCM1-FKBP and Kif5b-FRB in the presence of rapamycin, GFP-positive particles were excluded from perinuclear region and localized to the cell periphery (Fig. 5H; middle panel). In cells co-expressing GFP-PCM1-FKBP and BICD2-FRB in the presence of rapamycin, GFP-positive particles were accumulated at the centrosome (Fig. 5H; right panel). Thus, it was verified that the rapamycin-dependent chemical dimerization-mediated manipulation of PCM1 localization worked well in our experimental paradigm.

We exposed DEX-synchronized NIH/3T3 cells to 500 nM rapamycin (Rapa) for 1 h at 23 h after synchronization, and subsequently immunostained PCNT at 30 h after synchronization or measured the length of primary cilia visualized by immunostaining at 42 h after synchronization (Fig. 5I). Endogenous PCNT showed centriolar satellite-like distribution at 30 hr after DEX synchronization in cells transfected with GFP-PCM1-FKBP alone along with 1-hr rapamycin treatment (Fig. 5J; left panel). In cells co-expressing GFP-PCM1-FKBP and Kif5b-FRB along with 1-hr rapamycin treatment, endogenous PCNT disappeared from the centriolar satellites (Fig. 5J; middle panel). In cells co-expressing GFP-PCM1-FKBP and BICD2-FRB along with 1-hr rapamycin treatment, endogenous PCNT also disappeared from centriolar satellites and strongly accumulated on the centrosomes (Fig. 5J; right panel). We thus succeeded in perturbing the transient PCNT increase in centriolar satellites at 30 h after DEX synchronization. Under this condition, we investigated the length of primary cilia at additional 12 h after chemical dimerization induction (Fig. 5K). The increase in primary cilium length observed at 42 h after DEX synchronization in cells expressing GFP-PCM1-FKBP alone was suppressed in cells co-expressing GFP-PCM1-FKBP and Kif5b-FRB or GFP-PCM1-FKBP and BICD2-FRB (2.59 ± 0.02 µm in GFP-PCM1-FKBP alone, 2.15 ± 0.03 µm in co-expressing with Kif5b-FRB, 2.21 ± 0.07 µm in co-expressing with BICD2-FRB) (Fig. 5K). Together, these data indicate that the PCNT localization to centriolar satellites contributes to circadian rhythm-dependent changes of primary cilium length.

During the formation of primary cilia, centriolar satellites are known to promote the recruitment of intraflagellar transport (IFT) proteins to the centrosome (Hall *et al*, 2023; Ishikawa & Marshall, 2011). We also examined whether intraflagellar transport (IFT) proteins increased/decreased in primary cilia with circadian rhythmic pattern. In this study, immunostaining for IFT88 and Arl13b or γ-tubulin were performed in NIH/3T3 cells synchronized with DEX treatment (Fig. EV2). Rhythmic oscillation was observed in the fluorescence intensity of IFT88 in Arl13b-positive primary cilia, with a maximum at 40 h after synchronization (Fig. EV2A and EV2B). In contrast, no significant changes were observed in the fluorescence intensity of IFT88 on γ-tubulin-positive centrosomes (Fig. EV2C). These results suggest that circadian oscillation of PCNT in centriolar satellites generate circadian rhythm of primary cilium length by regulating IFT proteins recruitment to primary cilia.

### Circadian rhythm of primary cilia length contributes to activation of the Hedgehog (Hh) signaling pathway by eternal stimuli

Primary cilia are known to sense extracellular SHH and contribute to the activation of Hh signaling pathway (Gerdes *et al*, 2009). We examined effects of circadian rhythms of primary cilium length on Hh signaling by measuring the expression of *Gli1* and *Ptch1* mRNA, which are expressed downstream of the Hh signaling pathway, in the presence of Smoothened (SMO)-agonist (SAG). Cells were exposed to 10 nM SAG for 4 h when the primary cilium length was minimal (28 to 32 h after synchronization) or maximal (40 to 44 h after synchronization) (Fig. 6A). mRNA levels of *Gli1* and *Ptch1* were measured by qPCR. In cells not exposed to SAG, no significant difference in *Gli1* and *Ptch1* mRNA levels was observed between cells with the minimal and maximal primary cilium length (Fig. 6B). In cells exposed to 10 nM SAG, *Gli1* and *Ptch1* mRNA levels were more markedly increased when cells were exposed to SAG with primary cilium length minimal (28 to 32 h after synchronization) than when cells were exposed to SAG with primary cilium length maximal (40 to 44 h after synchronization) (Fig. 6B). These results indicate that circadian oscillations of primary cilium length contribute to the generation of circadian rhythm in responsiveness to external stimuli such as SHH.

**Figure 6.**
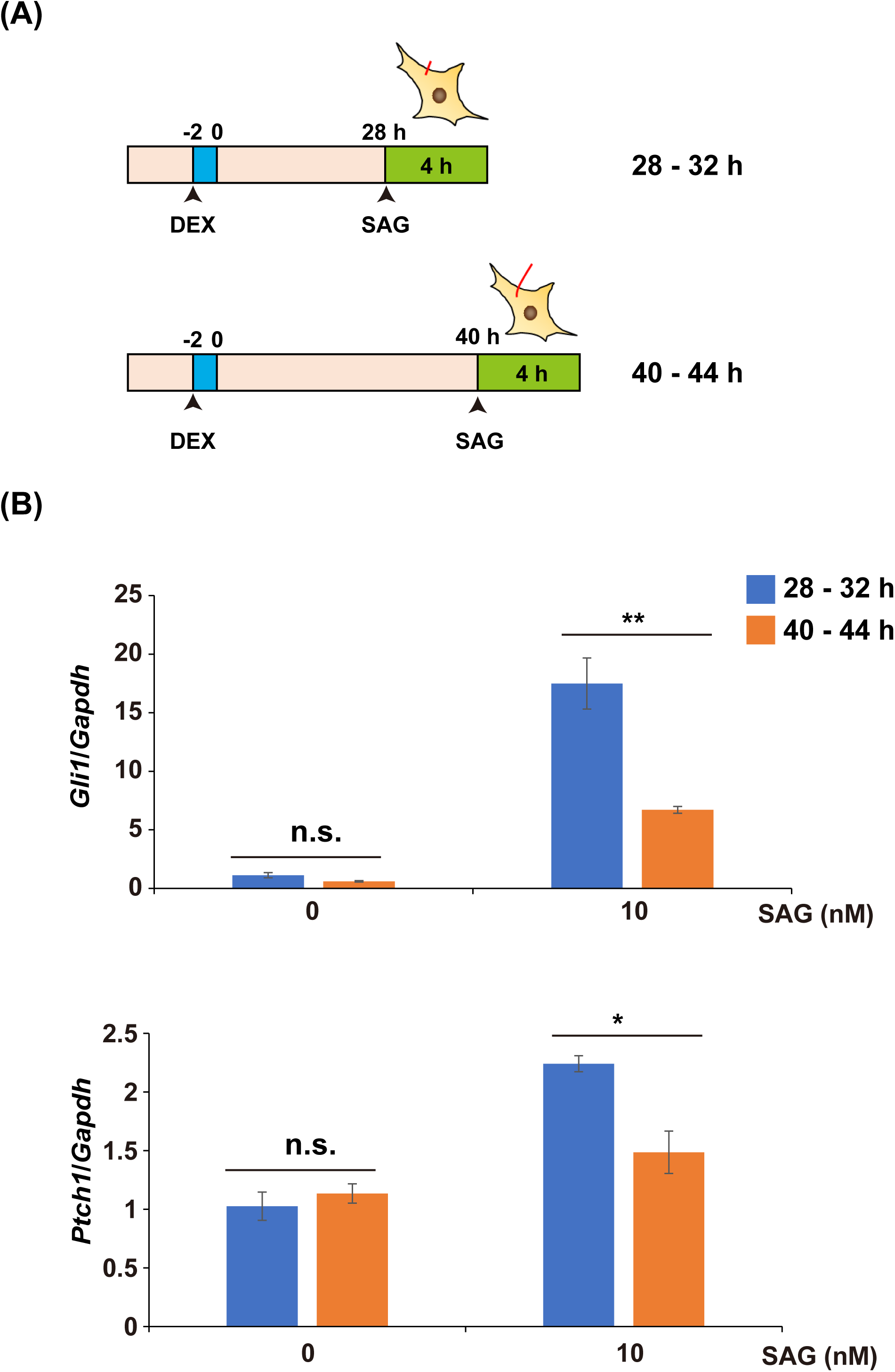
Rhythmic dynamics of primary cilium length influence SHH signaling. A) Schematic diagram of the protocol for treatment of SAG to NIH/3T3 cells exposed to DEX. B) Real-time qPCR analysis of *Gli1* or *Ptch1* mRNA expression levels in the NIH/3T3 cells incubated with SAG for 4 h at 28 h or 40 h after release from DEX. The data are from three independent experiments Data information: Data in panels B is presented as mean ± SEM. *P≤0.05 and **P≤0.01; n.s. indicates no significant difference (One-way ANOVA).

### Wound healing depends on the circadian phase of wounding, and its differences are eliminated by deficiency of primary cilia in NIH/3T3 cells

To investigate the physiological significance of circadian rhythms in primary cilium length, we performed a wound healing assay on synchronized NIH/3T3 cells. The cells were wounded when the primary cilium length was minimal (30 h after synchronization) or maximal (18 h or 42 h after synchronization), and the area of recovery was measured up to 24 h after wounding (Fig. 7A). We generated NIH/3T3 cells with deficient primary cilia, Kif3a knockout (KO) NIH/3T3 cells (Kif3a KO), as control references to evaluate the effect of primary cilia, and performed a wound healing assay with wild-type (WT) NIH/3T3 cells. The area of recovery in WT cells at 24 h after wounding was significantly larger for cells wounded when primary cilium length was minimal, rather than maximal (average recovery area 24.98 ± 0.66 µm^2^ at 30 h, 18.46 ± 1.56 µm^2^ at 18 h, and 16.16 ± 1.83 µm^2^ at 42 h) (Fig. 7B and 7C). In contrast, these differences were abolished by the loss of primary cilia in Kif3a KO (average recovery area 13.17 ± 2.56 µm^2^ at 30 h, 15.29 ± 2.89 µm^2^ at 18 h, and 8.39 ± 0.75 µm^2^ at 42 h) (Fig. 7B and 7C). We also generated Bbs2-KO NIH/3T3 cells (Bbs2 KO), which did not completely lose primary cilia but had very short primary cilia (Fig. EV3A). Similar to Kif3a KO, the changes of recovery ability after wounding upon the phase of cilia length were abolished in Bbs2 KO (Fig. EV3B and EV3C).

**Figure 7.**
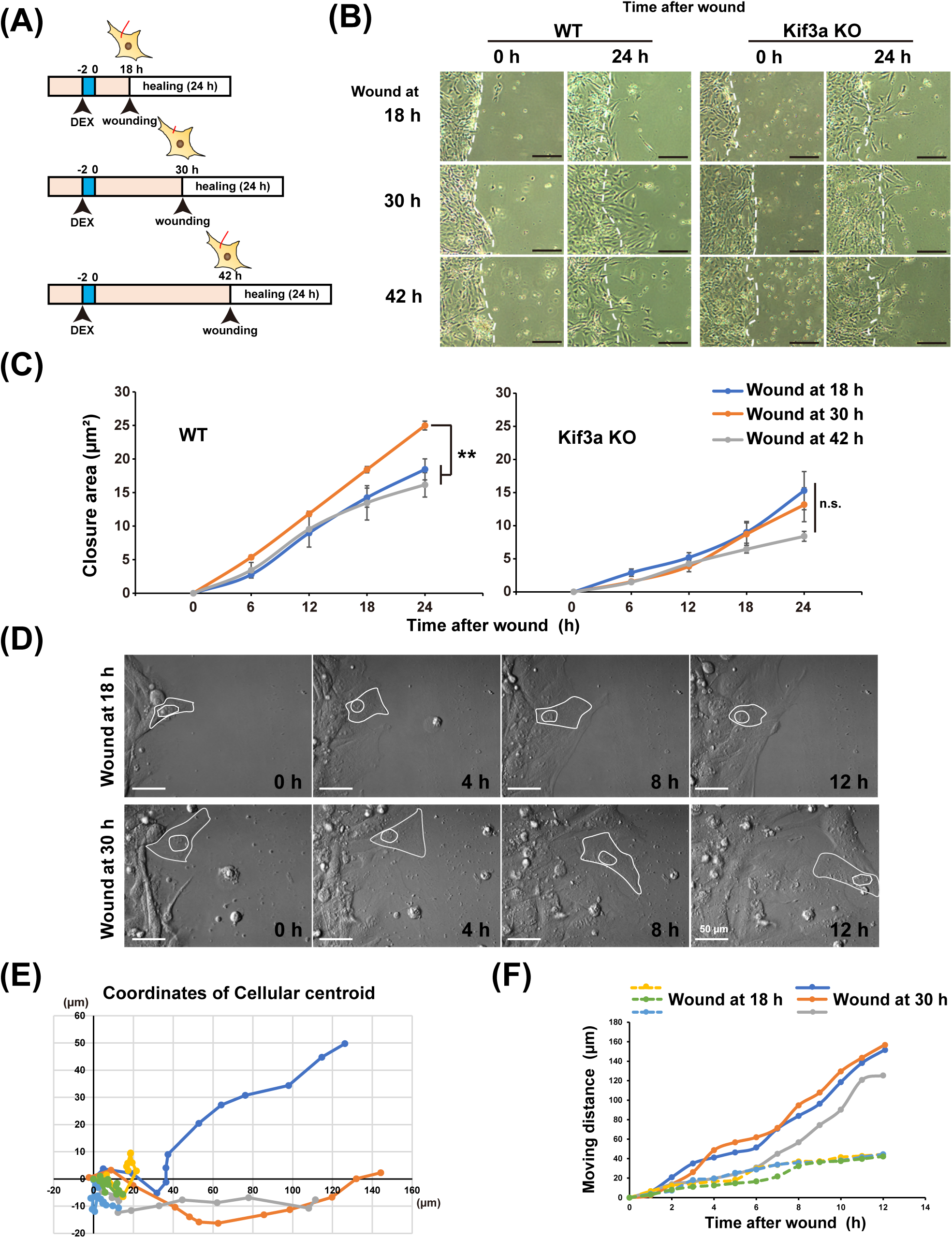
Primary cilia influence fibroblast mobilization with circadian changes in wound healing. A) Schematic diagram of the protocol for wound healing assay using NIH/3T3 cells exposed to DEX. B) Representative images of NIH/3T3 WT and Kif3a KO 0 or 24 h after wounding in the wound healing assay with the protocol illustrated in panel A. Scale bar, 200 µm. C) Quantitative analysis of the area of wound healing from panel B at the indicated time points after wounding. The data are from three independent experiments. Data are presented as the mean ± SEM. **P≤0.01; n.s. indicates no significant difference (One-way ANOVA). D) Representative time-lapse images of NIH/3T3 WT at the indicated time after wounding in the wound healing assay with the protocol illustrated in panel A. Scale bar, 50 µm. E and F) Quantitative analysis of the coordinates of the cellular centroid (E) and the moving distance of cells (F) from panel D.

We next examined more directly whether the length of primary cilia affected the capacity of healing after wounding. To this end, we compared cells that had normal primary cilia with cells that had long cilia. Long cilia cells were acquired by forced expression of Arl13b-venus, while normal cilia cells were produced by physiological expression of Arl13b-venus under endogenous Arl13b promoter (Fig. EV4A). Long cilia cells showed lower recovery ability after wounding than normal cilia cells (Fig. EV4B and EV4C). These results indicate that primary cilium length exhibiting circadian rhythms is involved in the circadian dynamics of cell motility.

To investigate in detail how the length of primary cilia exhibiting circadian rhythms affects cell motility, we performed time-lapse analysis. NIH/3T3 WT cells were wounded 18 or 30 h after synchronization and were photographed every 10 min. The coordinate migration and migration speed of these cells was determined. Cells with wounds at the time when the primary cilia length is minimal (30 h after synchronization) migrated almost linearly toward the wound (Fig. 7D, lower panels, 7E, and Movie EV1). In contrast, cells with wounds at the time when the primary cilia length was maximal (18 h after synchronization) did not move much and movement was not toward the wound (Fig. 7D, upper panels, 7E, and Movie EV2). The migration rate was also faster for cells wounded 30 h after synchronization than at 18 h (Fig. 7F). These results indicate that NIH/3T3 cells with shorter primary cilium lengths migrate faster than cells with longer primary cilium lengths and move in a straight line in the direction of the wound, leading to faster wound healing.

### Rhythmic oscillations of primary cilium length of NIH/3T3 cells toward wound repair is arrested

To further assess primary cilia in the wound healing assay, wounds were created in lawns of NIH/3T3 cell growth 18 or 30 h after synchronization, followed by immunostaining 12 h after wounding (Fig. 8A). At 12 h after wounding, three cell populations were observed based on differences in cell density: cells not facing the wound, cells facing the wound, and cells invading the wound area (Fig. 8B). The primary cilium length in NIH/3T3 cells wounded 30 h after synchronization was longer in cells facing the wound than in cells wounded at 18 h. The results were similar to the comparative lengths of primary cilia 42 h and 30 h after synchronization (average length: 3.67 ± 0.11 µm versus 2.27 ± 0.06 µm) (Fig. 8C and 8D). In cells wounded 30 h after synchronization, primary cilia in cells facing the wound were shorter (average length 2.31 ± 0.09 µm) than those in cells not facing the wound (Fig. 8C and 8D). The length of primary cilia in cells wounded 18 h after synchronization were a mixture of longer and shorter cilia in cells facing the wound (average length 3.18 ± 0.10 µm) (Fig. 8C and 8D). In cells invading the wound, primary cilia in cells wounded at 30 and 18 h after synchronization were absent or very short (Fig. 8C). These results indicate that the circadian rhythm of primary cilium length is maintained in cells not facing the wound, similar to the results of Figure 1E. In contrast, in cells facing the wound, the length of primary cilia was maintained at the time of wounding, indicating a lack of connection with the circadian rhythm. Furthermore, the absence or very short primary cilia in cells invading the wound area indicates that resorption of primary cilia is necessary for cell migration to repair the wound. Thus, wound healing could have been faster in cells wounded 30 h after synchronization than in those wounded 42 h after synchronization.

**Figure 8.**
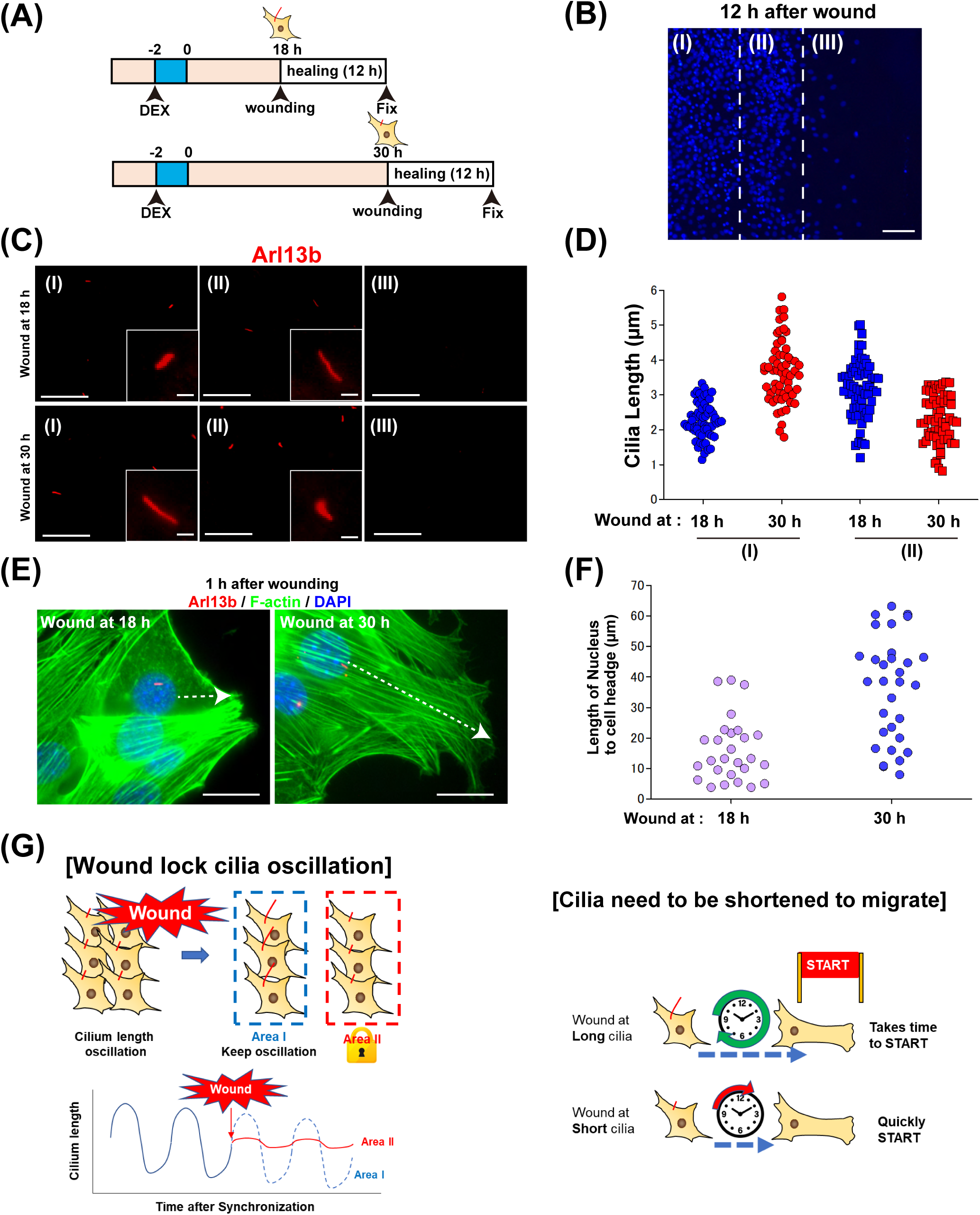
Rhythmic dynamics of primary cilia in wound healing. A) Schematic diagram of experimental protocol after the wound healing assay for NIH/3T3 cells exposed to DEX. B) Representative fluorescence image of DAPI-stained NIH/3T3 12 h after wounding using the protocol illustrated in panel A. Scale bar, 200 µm. C) Immunostaining for primary cilia (red; Arl13b) in NIH/3T3 cells at the indicated area (I-III as in B) with the protocol illustrated in panel A. Scale bar, 20 µm and 2 µm in magnificent images. D) Quantitative analysis of primary cilium length from panel C. The data are from over 60 cells per condition from two individual experiments. E) Immunostaining for primary cilia (red; Arl13b), F-actin (green; phalloidin), and DNA (blue; DAPI) in NIH/3T3 cells 1 h after wounding with the protocol illustrated in panel A. Arrows indicate the direction and length between the nucleus and the migrating edge of cell. Scale bar, 20 µm. F) Quantitative analysis of the distance from the nucleus to the migrating edge of cell from panel E (white arrows in panel E). The data are from 27 cells per condition. G) Hypothetical schematic diagram of the relationship between primary cilia and cell migration showing circadian rhythms.

To further investigate the effect of primary cilium length on cell migration in the wound healing assay, we analyzed the structure of actin filaments by fluorescent staining of F-actin with phalloidin. One hour after wounding, no significant change in actin polymerization was observed at either 18 or 30 h after synchronization (Fig. 8E). However, the distance from the cell nucleus to the tip of the cell in the direction of the wound was greater at 30 h than at 18 h after synchronization (Fig. 8E and 8F). No significant change was observed in the angle of the direction of the wound relative to the orientation of the primary cilia among these cells (Fig. 8E and Fig. EV5). These results indicate that short primary cilia can facilitate cell body changes in the direction of migration in the wound healing assay.

The collective findings of the wound healing assay can be summarized as follows. When a wound is introduced in a monolayer of cells, circadian changes in the length of primary cilia in cells contacting the wound are arrested. In addition, cell morphology changes in response to the wound and cell migration increase progressively as cilia length decreases. These observations suggest that wounds heal faster at the shortest cilia phase (30 h after synchronization) than at the longest cilia phase (18 or 42 h after synchronization) (Fig. 8G).

## Discussion

The most important finding of this study is that primary cilium length exhibits an oscillatory rhythm that coordinates with rhythmic clock gene expression. Ciliary structure regulates cell cycle progression (Plotnikova *et al*., 2009). Several clock genes act as gatekeepers in the cell cycle (Matsuo *et al*, 2003). BMAL1/CLOCK controls the cell cycle by directly regulating the expression of WEE1, an inhibitor of CDC2/cyclin B1 (Farshadi *et al*, 2019; Obi-Ioka *et al*, 2013). PER1 inhibits Cyclin D1 (Fu *et al*, 2016). In this study, we performed all experiments under conditions in which the concentration of serum in the culture medium was reduced from 10% to 1% to induce ciliogenesis by arresting the cells in the G0 phase of the cell cycle. Circadian rhythms in primary cilium length have also been observed in neurons, which are non-dividing cells. The circadian clock oscillates independently of the cell cycle (Matsuo *et al*., 2003). The formation of centrioles, which play a major role during cell division as the center of spindle formation, also exhibit a circadian rhythm independent of the cell cycle (Aydogan *et al*., 2020). Our current findings indicate that circadian rhythm or primary cilium length, regulated by clock genes, is a phenomenon that is independent of the cell cycle.

In this study, we observed oscillatory rhythms in the primary cilium length in cultured cells and in the murine brain. A previous study reported a relationship between circadian rhythms and ciliary protein expression in the brain (Baldi *et al*, 2021). In addition, it was reported that primary cilia in the SCN indicate circadian rhythms (Tu *et al*., 2023). Further analysis is needed to elucidate whether the mechanism of primary cilia *in vivo*, as in cultured cells, is mediated by the regulation of PCNT localization to centriolar satellites by clock genes. Whether primary cilia also exhibit circadian oscillations in tissues other than the brain, such as the kidneys, requires further investigation.

Immunostaining data in this study showed that approximately 50% of desynchronized NIH/3T3 cells were positive for primary cilia and 30-70% of synchronized NIH/3T3 cells were positive for primary cilia. Fraction of ciliated cells data are consistent with only 30-70% of NIH3T3 cells having detectable circadian regulation of primary cilia. These results indicate that circadian rhythms of primary cilium length observed in this study could be observed only in some population of NIH/3T3 cells. Live cell imaging using NIH/3T3 cells expressing Arl13b-vensu allowed us to track individual primary cilia for 72 hours, and we observed oscillations in primary cilium length with a cycle of approximately 24 hours. In addition, some cells were observed in which the fluorescent primary cilia disappeared or became apparent during the time-lapse imaging. These results suggest that primary cilia-positive cells in immunostaining are not always positive for 48 hours, and that primary cilia could shorten until they become undetectable. Therefore, circadian rhythm of primary cilium length observed in this study is observed in several cells.

SR9011 is a synthetic agonist of REV-ERB, the nuclear receptor involved in the cell-autonomous circadian transcriptional feedback loop as a transcriptional repressor. REV-ERB alters circadian behavior and the circadian pattern of core clock gene expression in the hypothalamus of mice (Solt *et al*, 2012). SR9011 applied at 20 µM is shown to inhibit autophagy in cancer cells (Sulli *et al*., 2018). The functional interaction between primary cilia and autophagy, and the autophagy-induced degradation of proteins essential for ciliogenesis have been described (Pampliega *et al*., 2013). In this study, the elongation of primary cilia by high concentrations of SR9011 thus could result from both inhibition of clock genes and blockade of the autophagy process. To eliminate the possibility of autophagy involvement, we tried lower concentrations of SR9011 and found that Bmal1 mRNA expression oscillations were suppressed, even with 2 µM SR9011, especially at 30 h after synchronization. SR9011 at 2 µM did not induce excessive elongation of primary cilia. Thus, the low concentration of SR9011 could inhibit circadian oscillations of primary cilia length by inhibiting only the expression of clock genes, not autophagy. In addition, the circadian rhythm of primary cilia length was abolished in Bmal1-KO cells, and the length of primary cilia was always the same as that of the WT cells at the time of the maximum length (40 to 44 h after synchronization). This result suggests that Bmal1 itself could function in an inhibitory manner on the formation of primary cilia. The transient increase in PCNT at the centriolar satellites 30 h after synchronization was also inhibited with 2 µM SR9011. These results suggest that oscillations in clock gene expression produce circadian rhythms in the length of primary cilia through direct regulation of PCNT expression in centriolar satellites. Further detailed analysis is required to confirm this suggestion.

PCNT is a component of centriolar satellites and contributes to centrosome stability (Quarantotti *et al*., 2019). PCNT forms a complex with intraflagellar transport (IFT) proteins and polycystin-2 (PC2), which is reportedly required for the assembly of primary cilia (Jurczyk *et al*., 2004). In contrast, in trisomy 21, the most prevalent human chromosomal disorder, an excessive increase in PCNT inhibits the transport of centrosome and cilia proteins, resulting in defects in cilia formation (Galati *et al*, 2018; Jewett *et al*, 2023; McCurdy *et al*, 2022). In this study, PCNT-positive centriolar satellites transiently increased in NIH/3T3 cells 30 h after synchronization when the primary cilium length was minimal. In addition, inhibition of microtubule polymerization with colchicine suppressed the transient increase in PCNT-positive centriolar satellites and significantly suppressed the primary cilium length maxima observed in NIH/3T3 cells 42 h after synchronization. Serum starvation-induced ciliogenesis is considered to begin within minutes of the onset of serum starvation, but the length of primary cilia becomes maximal 24 to 48 hours after the onset of serum starvation, indicating a time lag (Kukic *et al*, 2016; Westlake *et al*., 2011; Zhang *et al*, 2020). The increase in the number of ciliated cells by serum starvation is suppressed in centriolar satellites-deficient cells compared to normal cells, but the difference is observed at 18 h after the onset of serum starvation (Aydin *et al*., 2020). Although it is not reported how long it takes for the primary cilia to reach their maximum length from the formation of centriolar satellites, a time lag of about 12 h, as in the results of this study, is expected to exist. In this study, the amount of IFT88 in the primary cilia showed circadian oscillations in NIH/3T3 synchronized by DEX, which gradually increased from 28-32 hours after synchronization, reaching a maximum at 44 hours. Thus, it is likely that centriolar satellites causes slow recruitment of ciliary proteins to primary cilia, which would result in a time lag between the formation of centriolar satellites and the reaching of the primary cilium length maxima. our observations indicate that clock genes confer circadian rhythms on primary cilium length through the subcellular localization of PCNT-positive centriolar satellites. Further experiments should be performed to reveal the mechanisms by which clock genes regulate the subcellular localization of PCNT-positive centriolar satellites.

In this study, PCM1 subcellular localization showed no circadian rhythm, and a constant amount of PCM1 was always present in centriolar satellites. In addition, forced translocation of intracellular PCM1 using chemical dimerization system FKBP/FRB abolished distribution of PCNT-positive particles in centriolar satellites, and suppressed primary cilium length maxima in NIH/3T3 cells at 42 h after synchronization. PCM1 is known to serve as centriolar satellites scaffold protein that interacts with various proteins constituting the centriolar satellites (Jurczyk *et al*., 2004; Quarantotti *et al*., 2019). PCM1 in centriolar satellites does not show circadian rhythm, but could be essential for circadian rhythm of PCNT accumulation in centriolar satellites. Indeed, the subcellular localization of PCM1 and PCNT does not necessarily coincide (Guo *et al*, 2006; Staples *et al*, 2012). It has been reported that only about 48% of PCM1 and PCNT particles are in the same location (Kubo & Tsukita, 2003). PCM1 is required for PCNT to be present in the centriolar satellites, but PCNT is not necessarily present where PCM1 is present. Thus, PCNT puncta is not likely to be present in the same ectopic PCM1 puncta locations found in Kif5b-FRB-expressing cells.

The physiological significance of the circadian rhythm of primary cilium length requires further investigation. It is possible that changes in the length of primary cilia could produce circadian rhythms in their sensitivity as antennae to extracellular signals. In this study, both *Gli1* and *Ptch1* mRNA levels relative to *Gapdh* mRNA were better responsive to SAG at the time when primary cilia length is minimum (28-32 h after release from DEX), while the responses to SAG were poor at the time when primary cilia length is maximum (40-42 h after release from DEX). The result is well consistent with the work of that the upregulation of *Gli1* and *Ptch1* expression by SAG exposure is suppressed in cells with elongated primary cilia (Yoshida *et al*, 2020). Primary cilia reportedly play important roles in development. However, circadian rhythms in the SCN have not been observed in prenatal mice or rats (Sumová *et al*, 2006). Oscillations in clock gene expression are also not observed in embryonic stem cells and occur with cell differentiation (Yagita *et al*, 2010). Therefore, it is necessary to focus on primary cilia and the developmental stages of immature cells to elucidate the physiological significance of primary cilia that exhibit circadian rhythms.

A previous study reported that the cellular clock in fibroblasts modulates cell migration, which affects the efficacy of wound healing (Hoyle *et al*., 2017). Our observations from the wound healing assay also suggest a circadian rhythm in the mobilization of NIH/3T3 cells. In addition, the circadian rhythm of cell migration is absent in NIH/3T3 cells that have lost (Kif3a-KO) and denature (Bbs2-KO) their primary cilia, suggesting that rhythmic cell mobilization depends on the circadian rhythm of primary cilium length. Previous studies have reported that primary cilia regulate cell migration as antennae by detecting chemokines, such as platelet-derived growth factor (Christensen *et al*, 2013; Schneider *et al*, 2010). Previous studies have also reported that the tips of primary cilia orient in the direction of travel of the migrating cell (Albrecht-Buehler, 1977). In this study, no correlation was observed between the orientation of primary cilia and the direction of the wound site at 1 h in the wound healing assay. Other authors reported that primary cilia regulate neural migration through centrosomal dynamics (Tsai & Gleeson, 2005). Neural migration is essential for centrosome movement toward the leading edge and cell nuclei. AC3 in primary cilia produces cAMP, which activates protein kinase A to regulate centrosome dynamics and subsequent nuclear movement (Stoufflet *et al*, 2020). In this study, we observed that NIH/3T3 cells with short primary cilia migrated faster and in a straight line into wounds than did cells with long primary cilia, resulting in faster wound repair. Previous studies have not reported a relationship between primary cilium length and cell mobilization.

In the wound healing assay that used cells synchronized by DEX, circadian rhythms of primary cilium length were maintained in cells not facing the wound. In contrast, the length of primary cilia in cells facing the wound was not associated with the circadian rhythm and maintained their length at the time of wounding (Fig. 8G). A previous study reported that the rhythmic oscillation of clock gene expression depends on cell density and is altered in conditions of low cell density (Noguchi *et al*, 2013). Cell-to-cell communication has also been suggested to affect circadian rhythms (Fuhr *et al*, 2019). In addition, in the present study, cells that invaded wounds were devoid of primary cilia or had very short primary cilia, suggesting that cells migrating to wounds have short primary cilia. These findings further suggest that wound healing is faster in cells that are wounded when primary cilia are short, because the primary cilia remain short and fixed, allowing for rapid conversion to a cell type that can invade the wound (i.e., cells with no or very short primary cilia). In contrast, cells that are wounded when primary cilia are long take longer to convert to the cell type that invades the wound. Thus, cells facing the wound have a mixture of long and short primary cilia, and the wound appears to heal more slowly (Fig. 8G). Further experiments are needed to determine why cells that invade wounds must eliminate or shorten their primary cilia.

We hypothesize that primary cilia anchor for the centrosome. During cell division, disassembly of primary cilia could free the centrosome and allow it to move dynamically. This phenomenon is thought to occur during cell migration. Indeed, in this study, cells in the wound area had no or very short primary cilia. Our prediction from the results we obtained is that the centrosome can be easily released from short primary cilia more than from long primary cilia, and that centrosome dynamics and cell migration can commence sooner (Fig. 8G).

In conclusion, the autonomous cellular clock provides rhythmic oscillations in primary cilium length by regulating PCNT accumulation in centriolar satellites. Primary cilia contribute to the circadian dynamics of sensitivity to extracellular signals such as SHH and cell migration during wound repair. Our findings contribute to an improved understanding of the functional role of the non-SCN autonomous cellular clock system and primary cilium length alterations in a variety of physiological and pathological processes involving cell migration in wound repair and ciliopathies. Thus, clock genes may represent novel targets for the discovery and development of therapies for ciliopathies.

## Materials and Methods

### Cell culture and treatments

NIH/3T3 cells purchased from American Type Culture Collection (CRL-1658) were cultured in Dulbecco’s Modified Eagle’s Medium (DMEM)-High Glucose (044-29765, Wako) supplemented with 10% fetal bovine serum (FBS; Gibco) at 37°C in an atmosphere of 5% CO_2_. The workflow for the induction of ciliogenesis and clock gene synchronization is shown in Fig. 1. Ciliogenesis was induced by changing the culture medium to DMEM with 1% FBS for 24 h. Next, the cells were exposed to DEX (Sigma-Aldrich) for 2 h to synchronize the expression of clock genes. DEX was then washed out and various experiments were performed. SR9011 (Sigma-Aldrich) was added to the culture medium simultaneously with DEX and continued to be added to the medium after DEX had been washed out. Colchicine (Sigma-Aldrich) was added to the culture medium 28 h after synchronization, incubated for 2 h, and then washed out. Smoothened Agonist (SAG, Funakoshi) was added to the culture medium at 28 or 40 h after synchronization, incubated for 4 h, and then total RNA were extracted from cells.

### Generation of Knock-out cell line

Detailed methods for the generation of Knock-out (KO) cell lines have been previously described (Ijaz & Ikegami, 2021; Ijaz *et al*, 2022; Phua *et al*., 2017). Bmal1-KO, Kif3a-KO, and Bbs2-KO cells were generated using CRISPR/Cas9-based genome-editing techniques. To construct an all-in-one expression plasmid for KO, a 20-bp target sequence (Bmal1:TTGTCGTAGGATGTGACCGA, Kif3a:CTATAGACAGGCCGTCAGCG, Bbs2:CTGCAGGGTTTCGATCACGA) was subcloned into the U6-gRNA/CMV-Cas9-2A-GFP plasmid backbone (ATUM). Cells were plated at a density of 2 × 10^4^ per cm^2^ and were transfected with the all-in-one plasmid U6-gRNA/CMV-Cas9-2A-GFP using polyethylenimine. Three days after transfection, green fluorescent protein (GFP)-positive cells were sorted by fluorescence-activated cell sorting (FACS) using a FACS Aria2 SORP cell sorter (Becton Dickinson Biosciences). Single cell clones were generated by the limiting dilution culture method. Mutant cells were screened by double-strand break site-targeted PCR genotyping (Ijaz *et al*., 2022). KO cells were identified by DNA sequencing.

### Real-time qPCR

Total RNA was extracted from cells and reverse-transcribed to cDNA using the PrimeScript 1^st^ Strand cDNA Synthesis Kit (TaKaRa Bio). cDNA was used as a template for real-time qPCR analysis on CFX96 Touch (Bio-Rad) using specific primers targeting *glyceraldehyde 3-phosphate dehydrogenase* (*Gapdh*; 5’-AGGTCGGTGTGAACGGATTTG-3’ as the forward primer and 5’-TGTAGACCATGTAGTTGAGGTCA-3’ as the reverse primer), *Arntl* (5’-TGACCCTCATGGAAGGTTAGAA-3’ as the forward primer and 5’-GGACATTGCATTGCATGTTGG-3’), *Gli1* (5’-CCAAGCCAACTTTATGTCAGGG -3’ as the forward primer and 5’-AGCCCGCTTCTTTGTTAATTTGA -3’ as the reverse primer), *Ptch1* (5’-GCCTTCGCTGTGGGATTAAAG -3’ as the forward primer and 5’-CTTCTCCTATCTTCTGACGGGT -3’ as the reverse primer), and TB Green Premix Ex Taq II (TaKaRa Bio). In each sample, expression levels of the *Arntl* (BMAL1 gene symbol), *Gli1*, *Ptch1* were normalized to *Gapdh* expression as an internal control.

### Mice

C57BL/6J mice purchased from Japan SLC, Inc. were maintained under conditions of controlled temperature and humidity with a 12-h light/dark cycle and *ad libitum* access to standard laboratory food and water. The experiments involving mice were approved by the Natural Science Center for Basic Research and Development at Hiroshima University.

### Immunocytochemistry

Cells plated on cover glass (MATSUNAMI) were fixed with 4% PFA for 15 min at 37°C or 100% MeOH for 10 min at 4°C. The cells were blocked with 5% normal goat serum containing 0.1% Triton X-100 in PBS (blocking buffer) for 1 h at room temperature (22 to 28°C). The primary antibodies were diluted in blocking buffer and incubated with cells overnight at 4°C. The cells were immunostained with anti-Arl13b (Protein Tech, 17711-1-AP or Abcam, ab136648; both 1:1000), anti-pericentrin (Abcam, ab220784; 1:400), anti-γ-tubulin (Sigma-Aldrich; GTU88, T6557; 1:400), anti-α-tubulin (Sigma-Aldrich, T9026; 1:400), anti-PCM1 (Protein Tech, 19856-1-AP; 1:400), anti-IFT88 (Protein Tech, 13967-1-AP; 1:400) and anti-GFP (MBL, 598; 1:400). The cells were washed with PBS to remove unbound antibody and incubated for 2 h with Alexa Fluor-conjugated secondary antibodies (Invitrogen; 1:500), 4′,6-diamidino-2-phenylindole (DAPI; Wako; 1:1000), and Alexa488-conjugated phalloidin (1:1000) at room temperature. After washing with PBS, the cover glasses were mounted on glass slides with VECTASHIELD mounting medium (Vector Laboratories). Images were acquired using a model DMI3000B epifluorescence microscope (Leica) and model FV1000D confocal microscope (Olympus). Quantitative image analyses were performed using ImageJ software (NIH).

### Immunohistochemistry

Adult 8–12 week old male mice were deeply anesthetized with isoflurane before intracardial perfusion with PBS and 4% PFA. The brains were post-fixed overnight and cryoprotected in 30% sucrose. The brains were embedded in OCT tissue TEC (Sakura) and 30 µm-thick frozen coronal sections were cut using a model CM 3050 cryostat (Leica). The sections were blocked with blocking buffer for 1 h and incubated with primary antibodies overnight at 4°C. The antibodies used for the brain sections in this study were anti-Arl13b (Abcam; 1:400), anti-ACIII (Novus, NBP1-92683SS; 1:1000), anti-NeuN (Abcam, ab104224; 1:400), and anti-GFAP (Abcam, ab4674; 1:400). After washing with PBS, the sections were incubated for 2 h with Alexa Fluor-conjugated secondary antibody (Invitrogen, 1:400) and DAPI (Wako, 1:1000). Confocal images were acquired using a model FV1000 microscope (Olympus).

### Western blotting

Cells were solubilized in 1× sodium dodecyl sulfate-polyacrylamide gel electrophoresis (SDS-PAGE) sample buffer (Wako). Lysate samples were subjected to 6% SDS-PAGE and transferred to polyvinylidene difluoride membranes. Membranes were blocked with 5% bovine serum albumin (BSA) in Tris buffered saline containing 0.1% Tween 20 (TBST) and immunoblotted using primary antibody to pericentrin (Abcam) diluted 1:1000 in 1% BSA in TBST overnight at 4°C. After washing with TBST, the membranes were incubated for 1 h with horseradish peroxidase-conjugated secondary antibody (1:5000) and developed using ECL Prime (GE).

### Primary cilia imaging in live cells

Generation of NIH/3T3 cells expressing Arl13b-Venus in a physiological manner have been previously described (Ijaz & Ikegami, 2021; Nakazato *et al*, 2023). NIH/3T3 cells expressing Arl13b-Venus were seeded in glass bottom dishes, fluorescent images were taken every hour using LCV110 (OLYMPUS), which is embedded in CO_2_ incubator, and equipped with a 40x objective lens (NA 0.95, UPlanSApo40X; Olympus) and a camera (EXi Aqua; Q Imaging). The images were taken by using a 2x2 binning mode, and the pixel size of images is 0.32 µm/pixel. The length of primary cilia was measured using Image J.

### Chemical dimerization assay

Plasmids for expressing GFP-PCM1-FKBP, HA-Kif5b (1-269 aa)-FRB, and HA-BICD2 (1-198 aa)-FRB were generous gifted from Dr. Karalar (Aydin *et al*., 2020). Cells were co-transfected with GFP-PCM1-FKBP and HA-Kif5b-FRB or HA-BICD2-FRB. One-day after transfection, culture medium was replaced with DMEM containing 1%FBS, and after an additional 24 h, the cells were synchronized by DEX. At 23 h after release from DEX, cells were treated with 500 nM rapamycin (Sigma) for 1 h. Six or eighteen hours after rapamycin induction, cells were fixed and stained with indicated antiboidies.

### Wound healing assay

A scratch was made in a cell monolayer using a sterile 200 µL plastic pipette tip at the indicated time after removing DEX. Images were acquired every 6 h for 24 h after wounding. Cell closure of the wound was quantitated using ImageJ software. Live cell imaging by time-lapse analysis was performed using a model LCV110 microscope (OLYMPUS) every 10 min for 12 h at 37°C. Quantification of the coordinates of the cellular centroid and cellular moving distance was performed using ImageJ software. In post-scratch immunostaining, cells invading the wound area is defined as cells that have invaded the wound and are not in contact with other cells, and cells facing the wound is defined as cells that are in contact with other cells but facing the wound.

### Quantification

Quantitative image analysis was performed using ImageJ software (NIH). Detailed methods for the measurement of the length of primary cilia have been previously described (Nakazato *et al*., 2023). Lengths of primary cilia are measured by tracing the primary cilia with the “Segmented Line” in ImageJ. Calculation of the length of primary cilia traced is performed by calculating the number of pixels per µm. The number of ciliated cells is calculated by counting the number of DAPI and Arl13b-positive cilia in 5-fields of view per sample (a total of at least 50 cells).

The puncta density of Pericentrin was measured according to a previous study (Galati *et al*., 2018). PCNT-positive particles were enclosed using “Polygon selections” and XY-coordinates of each were measured by the “Analyze particles” in ImageJ. Measure the distance from the center of centrosome to each particles and indicate the number of particles at the indicated distance.

### Statistical analysis

All experiments and quantitative analyses were blinded and randomized to remove bias. The results are presented as the mean ± SEM. Group means were compared using One-way ANOVA. Statistical significance was set at P < 0.05.

## Data availability

Our study includes no data deposited in public repositories.

## Acknowledgments

This work was supported in part by JSPS Grants-in-Aids for Young Scientists (20K15792) to RN and for Scientific Research (C) (21K06172) to KI, and Japan Science and Technology Agency, Precursory Research for Embryonic Science and Technology (JPMJPR17H1) to KI. Plasmids (peGFP-C1-PCM1-FKBP, pCI-neo-HA-Kif5b (1-807)-FRB, and pCI-neo-HA-BICD2-N (1-594)-FRB) were a kind gift from Dr. Elif Nur Firat-Karalar (Koç University). We thank Madoka Hamada, Mimoko Katoh, and Yoko Hayashi for technical assistance. A part of this work was the result of using research equipment shared in the Analysis Center of Life Science, Natural Science Center for Basic Research and Development, Hiroshima University, the Research Institute for Radiation Biology and Medicine, Hiroshima University (NBARD-00002) and the MEXT Project to promote public utilization of advanced research infrastructure (Program for supporting construction of core facilities; Grant Number JPMXS0441300023).

---

**Figure EV1.**
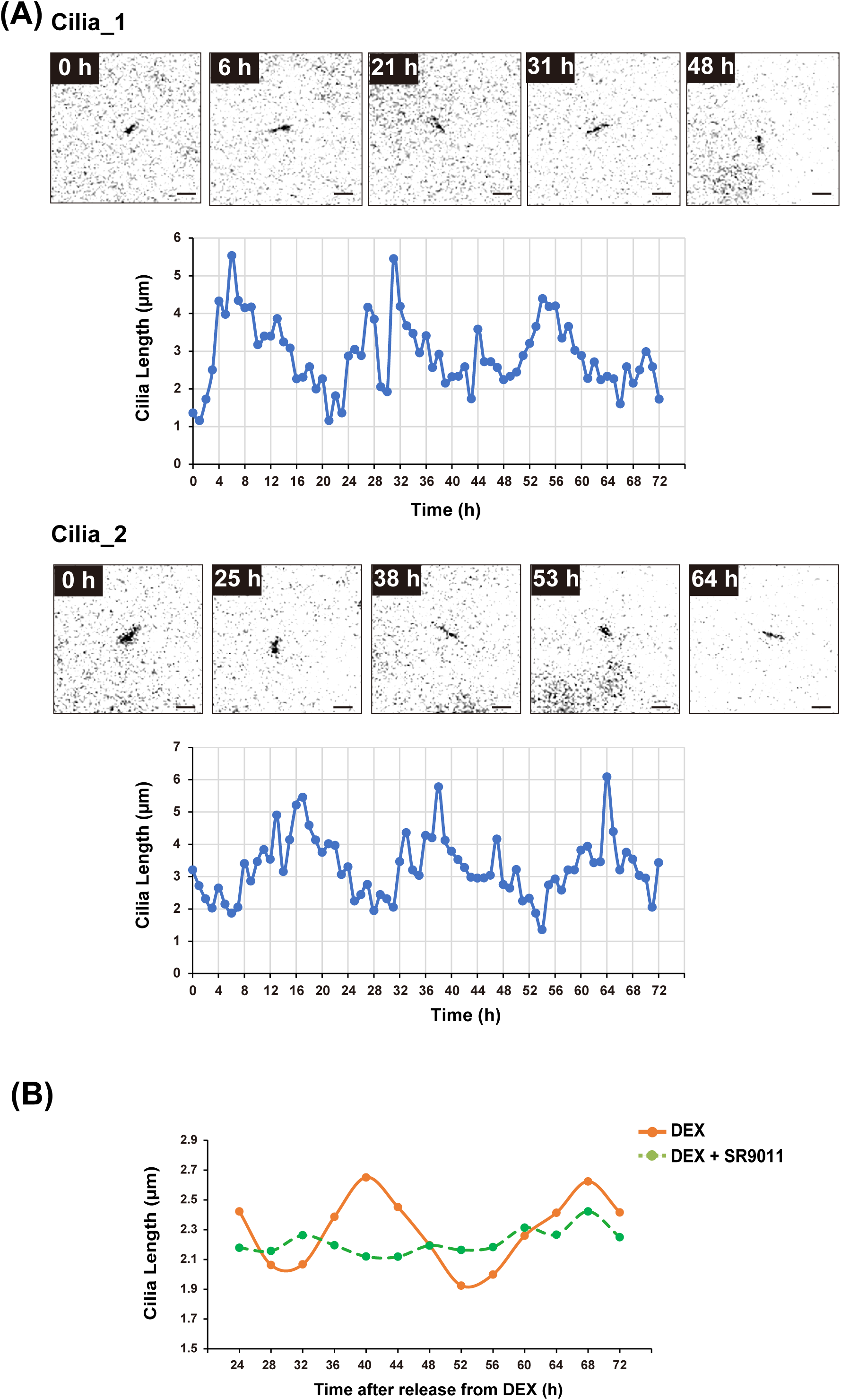
Circadian rhythm of primary cilium length is continuous for more than 48-hours. A) Representative time-lapse images of Arl13b-venus expressing NIH/3T3 at the indicated time and quantitative analysis of primary cilium length from Movie EV3 and EV4. Scale bar, 2 µm. B) Quantitative analysis of primary cilium length in NIH/3T3 cells treated with 2 µM SR9011 at each indicated time point after release from DEX.

**Figure EV2.**
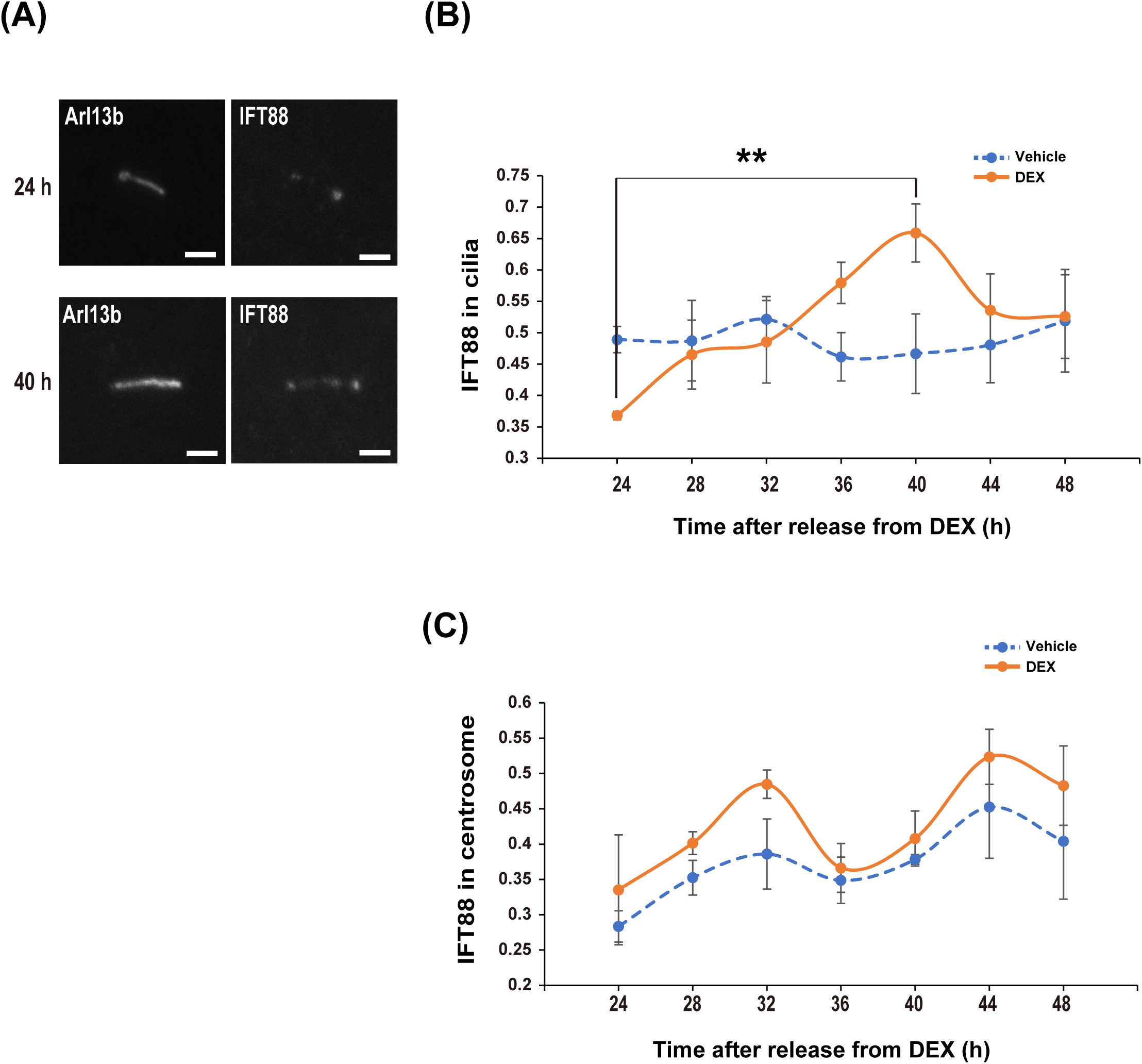
Rhythmic dynamics of IFT88 in primary cilia. A) Immunostaining for primary cilia (Arl13b) and IFT88 in NIH/3T3 cells at 24 or 40 h after release from DEX. Scale bar, 2 µm. B and C) Quantitative analysis of fluorescence intensity of IFT88 in primary cilia (Arl13b, B) and centrosome (γ-tubulin, C) of NIH/3T3 at the indicated time point after release from DEX. The data are from three independent experiments. Data information: Data in panels B and C are presented as mean ± SEM. **P≤0.01 (One-way ANOVA).

**Figure EV3.**
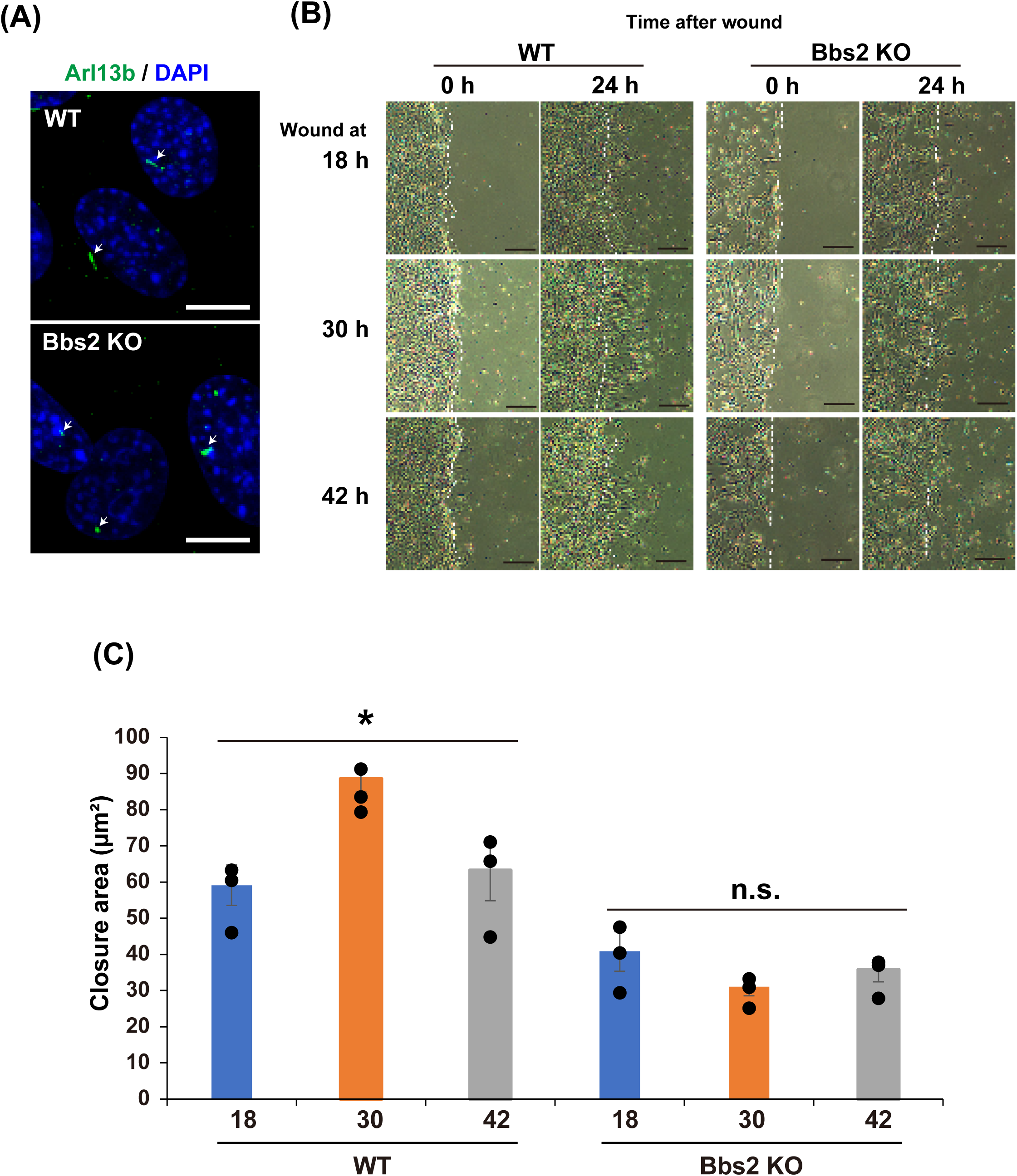
Bbs2-knock out NIH/3T3 in wound healing. A) Immunostaining for primary cilia (red; Arl13B) and DNA (blue; DAPI) in WT or Bbs2-KO NIH/3T3 cells. Scale bar, 10 µm. White arrows point to primary cilia. B) Representative images of NIH/3T3 WT and Bbs2-KO 0 or 24 h after wounding in the wound healing assay with the protocol illustrated in Figure 7 panel A. Scale bar, 200 µm. C) Quantitative analysis of the area of wound healing from panel B at 24 h after wounding. The data are from three independent experiments. Data information: Data in panels C is presented as mean ± SEM. *P≤0.05; n.s. indicates no significant difference (One-way ANOVA).

**Figure EV4.**
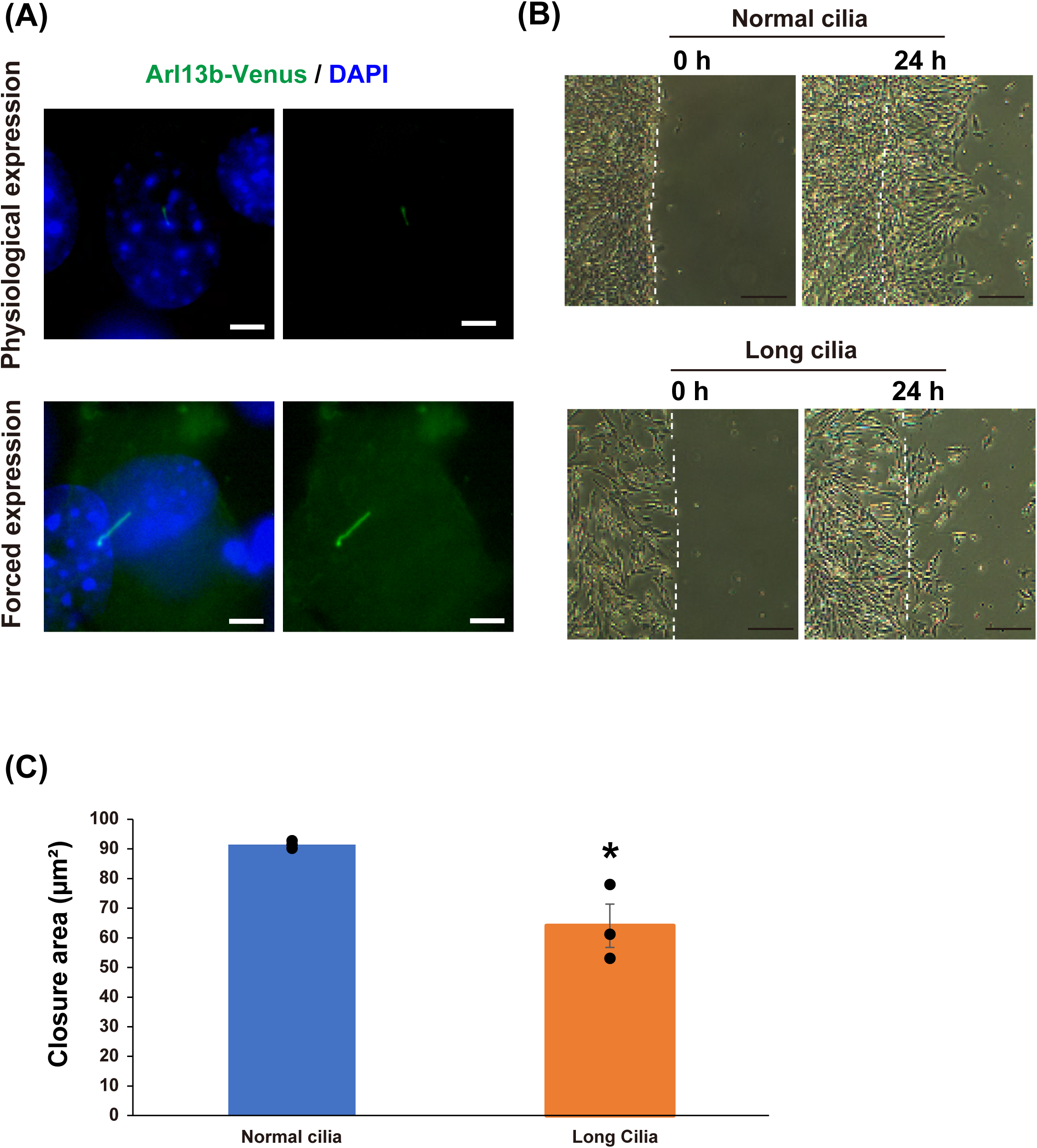
Increased primary cilium length decreases cell migration. A) Immunostaining for GFP in NIH/3T3 cells with forced expression of Arl13b-Venus (Long cilia) or physiological expression of Arl13b-Venus (Normal cilia) at 0 or 24 h after wounding in the wound healing assay. Scale bar, 5 µm B) Representative images of NIH/3T3 cells with long cilia or normal cilia at 0 or 24 h after wounding in the wound healing assay. Scale bar, 200 µm. C) Quantitative analysis of the area of wound healing from panel A at 24 h after wounding. The data are from three independent experiments. Data information: Data in panels C is presented as mean ± SEM. *P≤0.05 (One-way ANOVA).

**Figure EV5.**
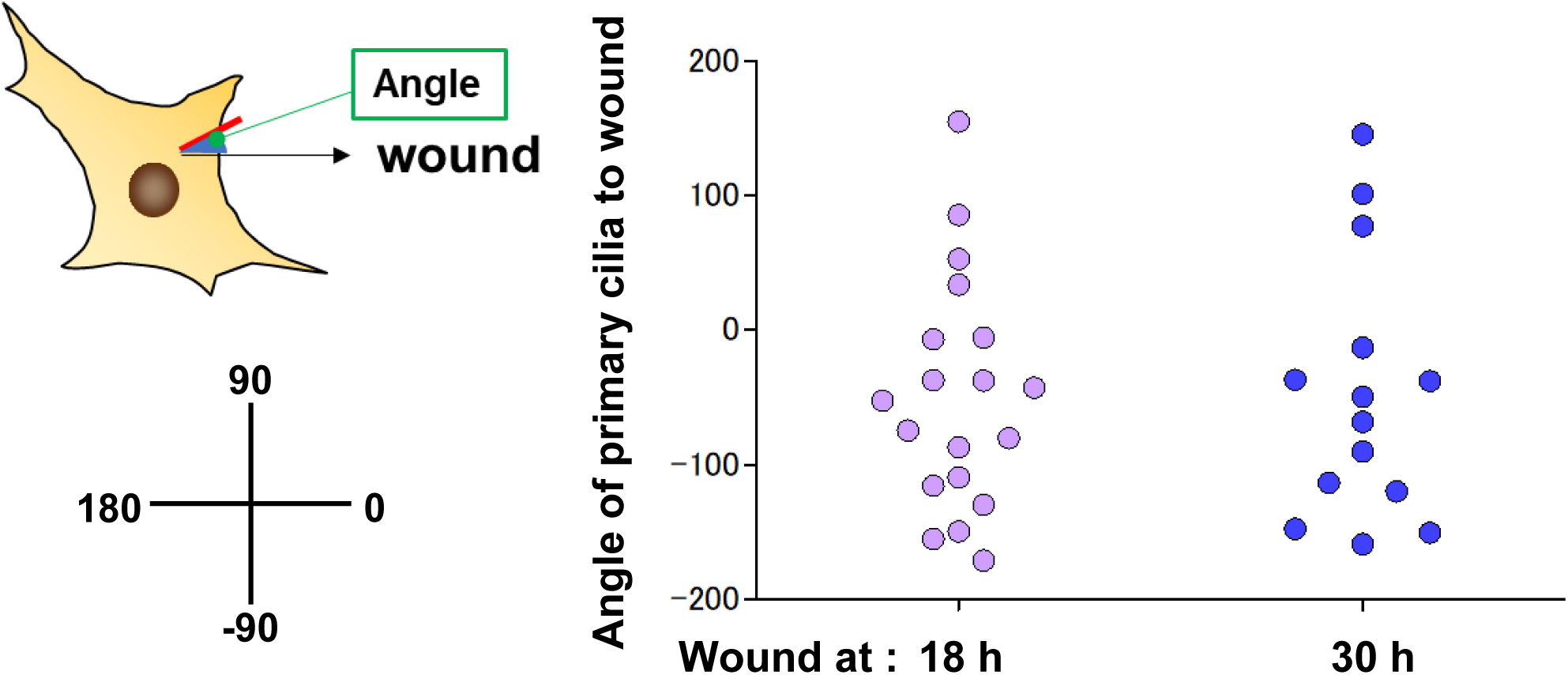
Primary cilia angle in wound healing. A) Schematic diagram of the angle of primary cilia relative to the direction of the wound. B) Quantitative analysis of the angle of primary cilia from panel E in Figure 8. The data are from 19 cells at 18 h and 14 cells at 30 h.

